# Spatiotemporal variation in habitat suitability predicts genomic diversity and structure in a Western Ghats endemic Tarantula (*Thrigmopoeus truculentus*)

**DOI:** 10.64898/2026.02.06.704511

**Authors:** Aritra Biswas, Praveen Karanth

## Abstract

Physical barriers are well known to restrict gene flow and generate population structure, yet what drives genetic differentiation in the absence of such barriers remains less understood. In these cases, long-term climatic fluctuations may shape genomic variation by altering habitat connectivity through time. The Western Ghats mountains in Peninsular India, marked by high endemism and *in-situ* radiations, provide a compelling natural laboratory to understand how historical climate change can shape genetic diversity. While the role of topographic barriers in generating diversity in this landscape is well documented, the influence of paleoclimatic processes has rarely been examined, especially from a genomic perspective. Here, we combine genome-wide SNPs and ecological niche modelling with present and past climatic data to test the role of elevation, geographic distance, environment and paleohabitat dynamics in shaping the genetic diversity and structure in a wet-adapted tarantula species endemic mostly to the central Western Ghats. Despite overall genomic admixture and continuity, populations show little north–south structuring, and genetically distinct central populations. Paleoclimate projections from the present to the Last Glacial Maximum reveal that northern and southern regions maintained stable habitat suitability for *Thrigmopoeus*, whereas central regions experienced high temporal variability. Linear mixed models identify historical stability of suitable habitats as the strongest predictor of genetic structure. Central populations occupying historically unstable habitats also show reduced heterozygosity, elevated inbreeding, and smaller historical effective population sizes. These results demonstrate that, even in the absence of physical barriers, long-term climate dynamics can generate and maintain fine-scale, within-species genetic diversity.

## 1. Introduction

Physical barriers such as mountain ranges, deep valleys and rivers are among the most well-known forces restricting gene flow and driving population differentiation (Fonseca et al., 2024; Y.-S. Li et al., 2019; Nazareno et al., 2019; Wang et al., 2025; Wei et al., 2013). When these barriers prevent dispersal, populations accumulate genetic divergence through mutation, drift, and often even through selection (Faria et al., 2021; Kulmuni et al., 2020; Schluter & Rieseberg, 2022). However, many species exhibit pronounced genetic structure even in the absence of conspicuous physical barriers to gene flow (Chaitanya et al., 2025; Hazzi & Wood, 2025; Hinckley et al., 2025). Studying processes that shape such phylogeographic patterns will contribute towards understanding the origin and maintenance of genetic variation. In such systems, population genetic structuring can arise from a combination of processes that operate without overt geographic isolation, including isolation by distance (spatial limitations to dispersal) (Cordeiro et al., 2019; Hardy & Vekemans, 1999; Pusadee et al., 2009), isolation by environment or climate (adaptive differentiation along environmental gradients) (Hu et al., 2022; Segovia et al., 2020), habitat fragmentation and microhabitat specialization (Day et al., 2025; Samad- & Sandra, 2023), as well as historical and paleoclimatic processes that have repeatedly altered connectivity through time (Batalha-filho et al., 2023; Ortego et al., 2023).

Among the suite of non-physical processes shaping population structure, growing evidence highlights paleoclimatic fluctuations as a key determinant of genetic differentiation through time (Fonseca et al., 2023; Morgan et al., 2011; Willis et al., 2004). Repeated cycles of glacial and interglacial periods during the Quaternary period have caused dramatic shifts in temperature and precipitation, leading to range contractions, expansions, and local extinctions across many taxa (de Pous et al., 2016; Fonseca et al., 2023; Lima et al., 2017; Morgan et al., 2011; Nogués-Bravo et al., 2010; Pearce-Higgins et al., 2015; Willis et al., 2004). Such climatic oscillations can alter patterns of habitat suitability and connectivity, thereby influencing gene flow even in the absence of geographic barriers (Hazzi & Wood, 2025; Maier et al., 2019; Zhang & Zhang, 2012). Populations that persisted in climatically stable refugia often retain higher genetic diversity, whereas those that experienced repeated episodes of isolation and reconnection in unstable regions tend to show reduced diversity, increased inbreeding, and stronger differentiation (de Pous et al., 2016; Fonseca et al., 2023; Lima et al., 2017; Willis et al., 2004). Thus, paleoclimatic dynamics could serve as temporal barriers to gene flow, fragmenting and reconnecting populations through time and therefore leaving genomic signatures of past environmental fluctuations. Although these processes are thought to be more pronounced in temperate regions (Biswas & Karanth, 2025; Janzen, 1967; Pelletier & Carstens, 2018; Zhang & Zhang, 2012), there is growing evidence from tropical regions, often considered climatically stable, that even relatively modest climatic changes can have substantial evolutionary consequences (Alves-Ferreira et al., 2025; Fonseca et al., 2024; Janzen, 1967; Kanagaraj et al., 2023; Lima et al., 2017; Oswald et al., 2022).

Stretching for over 1,600 km along the western coast of peninsular India (8°N to 21°N), the Western Ghats form one of the most distinctive tropical mountain systems and a globally recognized biodiversity hotspot (Dahanukar et al., 2004; Subramanyam & Nayar, 1974). Rising steeply from the coastal plains in the west and giving way to the drier Deccan Plateau to the east, the range encompasses sharp gradients in elevation, rainfall, and vegetation that together create a striking mosaic of habitats, from wet evergreen forests and montane sholas to semi-evergreen and dry deciduous formations (Davidar et al., 2007; Gopal et al., 2023; Subramanyam & Nayar, 1974). This environmental heterogeneity, shaped by complex topography and the orographic effects of the southwest monsoon, has promoted exceptional levels of endemism and in situ diversification across a wide array of taxa (Davidar et al., 2007; Gimaret-Carpentier et al., 2003; Joshi & Karanth, 2013; Subramanyam & Nayar, 1974). Paleoclimatic reconstructions suggest that while parts of the WG range have remained persistently wet over time, other areas have experienced episodic climatic instability (Singh et al., 2006). These characteristics make the Western Ghats an ideal system for understanding how long-term climate dynamics, acting upon a complex geography, have shaped the spatial distribution of genetic diversity in tropical species.

Geographically, the Western Ghats is interrupted by a few prominent low-elevation valleys in the mountain chain, most notably the Palghat Gap (∼11° N, 40 km wide) and the Shencottah Gap (∼9° N, 7.5 km wide) (Rajesh et al., 1998; Soman et al., 1990). The influence of these features on shaping patterns of genetic and species diversity has been extensively studied across multiple taxa (reviewed in Biswas & Praveen Karanth, 2021). In mammals (Kolipakam et al., 2019; Ram et al., 2015; Vidya et al., 2005) and birds (Robin et al., 2010, 2015), these gaps often correspond to pronounced within-species genetic structuring, whereas in amphibians (Vijayakumar et al., 2016), squamates (Chaitanya et al., 2019; Mallik et al., 2020), and several invertebrates (Joshi & Karanth, 2013; Klaus et al., 2014; Sekar & Karanth, 2013), they have been associated with deep phylogenetic splits and cladogenetic events. However, for many other taxa, these gaps do not appear to impede gene flow (check the references in Biswas & Praveen Karanth, 2021 for details), suggesting that physical barriers alone cannot fully explain the observed patterns of genetic differentiation in the region.

In contrast, the potential role of climatic heterogeneity and paleoclimatic dynamics in shaping genetic structure within species remains mostly unexplored. Some studies have highlighted the influence of long-term climatic stability and environmental gradients on interspecific diversity—for instance, southern Western Ghats refugia preserving ancient clades (Gopal et al., 2023; Joshi & Karanth, 2013), Pleistocene climate-driven diversification in frogs (Cyriac et al., 2024; Vijayakumar et al., 2016), and the role of climatic stability in determining woody plant richness (Page & Shanker, 2020). Yet, these studies primarily address broad, macroevolutionary patterns. Explicit genomic investigations linking paleoclimatic fluctuations to intraspecific genetic structure are notably lacking. Recently, Chaitanya et al. (2025) demonstrated that local adaptation to precipitation seasonality in concomitance with Pleistocene climatic oscillations likely shaped and maintained speciation in an endemic Flying lizard (*Draco dussumieri*), despite the absence of physical barriers to gene flow. Another phylogeographic study showed that black pepper (*Piper nigrum*) from the Western Ghats does not exhibit genetic structure corresponding to physical gaps; instead, its genetic patterns appear to have been shaped by postglacial climatic processes (Sen et al., 2025). Given this context, it is plausible that long-term climate dynamics, rather than geography, play a dominant role in shaping population connectivity and genetic diversity across the Western Ghats for species whose distributions do not span major gaps.

To investigate what shapes genetic variation, we focused on the endemic tarantula *Thrigmopoeus truculentus* (Family Theraphosidae, Order Araneae), a burrowing mygalomorph spider endemic to the Western Ghats (Figure 1). This species has a narrow and continuous distribution concentrated in the central Western Ghats sensu Subramanyam & Nayar (1974), extending marginally into the southern portion of the northern Western Ghats. Its known range spans from Amboli (∼16°), Maharashtra, in the north to Wayanad (∼11.5°), Kerala, in the south, encompassing the entire Western Ghats tract of the states of Karnataka and Goa (Jose, 2018; Siliwal & Molur, 2009). Within this range, *T. truculentus* is locally abundant and inhabits hill-forested regions without encountering any major topographic barriers to dispersal such as valleys or rivers (Siliwal & Molur, 2009). Given this continuity, the species provides a compelling system to test how genetic variation and population structure arise in the absence of obvious physical barriers to gene flow. Moreover, mygalomorph spiders usually exhibit sedentary lifestyles and limited dispersal— characteristics that can lead to genetic structuring even at short spatial scales (Monjaraz-Ruedas et al., 2024).

**Figure 1:**
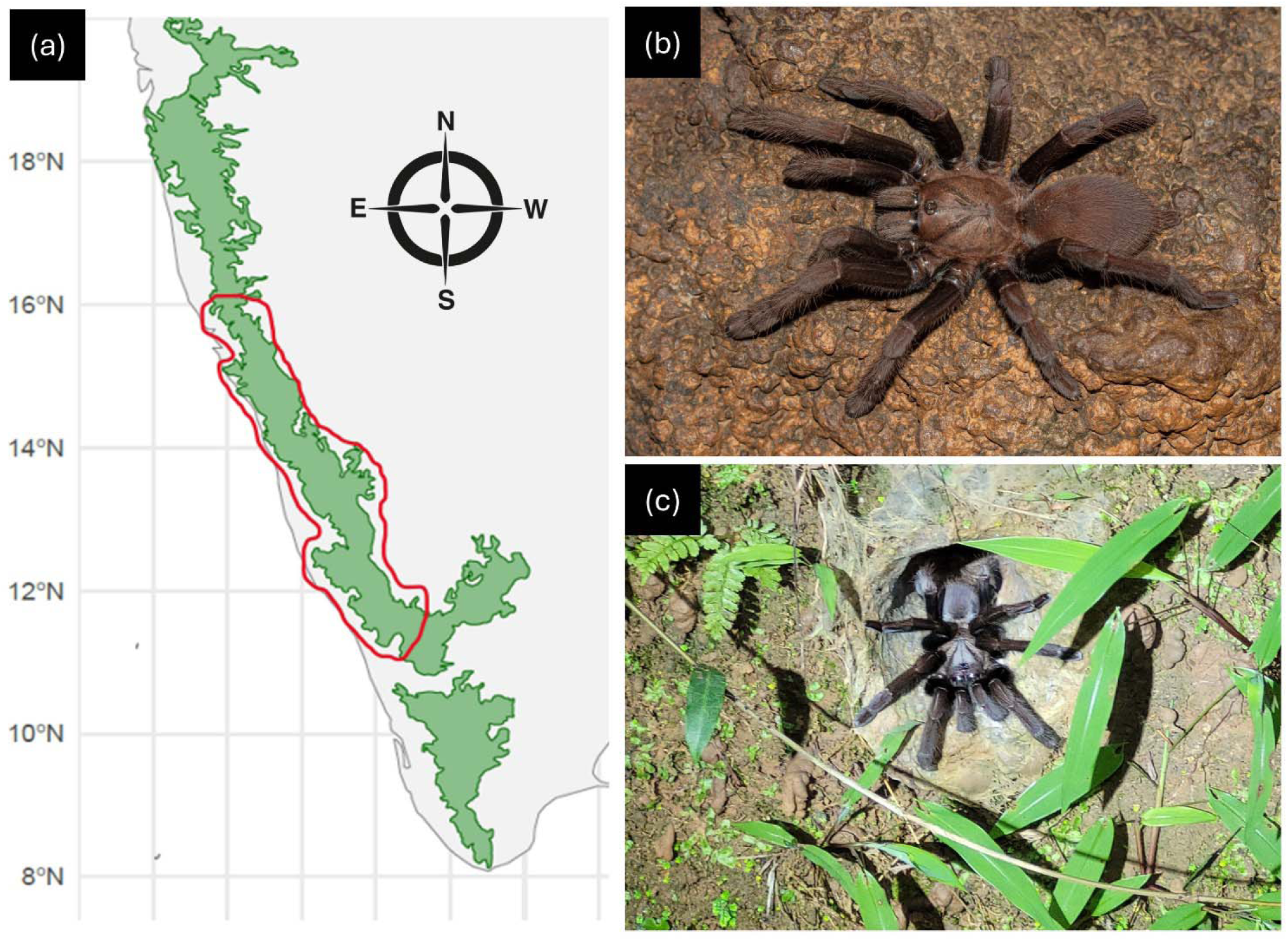
Distribution range of *Thrigmopoeus truculentus* and photos of the model organism in the wild.

We therefore evaluated four alternative, but not mutually exclusive, hypotheses to explain patterns of genetic structure and diversity in *T. truculentus*. First, genetic differentiation may arise through isolation by distance (IBD), whereby limited dispersal leads to increasing genetic divergence with geographic separation. Second, population connectivity may be constrained by topographic heterogeneity, with elevational differences acting as barriers to gene flow. Third, genetic structure may reflect isolation by environment, such that populations occupying habitats with differing suitability experience reduced connectivity. Finally, genetic differentiation may be primarily driven by long-term paleohabitat dynamics, where temporal variability in habitat suitability alters connectivity through time.

To test these competing hypotheses, we evaluated geographic distance, elevational differences (as a proxy for topographic heterogeneity), mean habitat suitability since the Last Glacial Maximum, and temporal variability in habitat suitability as predictors of genetic differentiation and diversity using a mixed-effects modelling framework. By explicitly contrasting these alternative mechanisms, we aim to disentangle the relative contributions of spatial, topographic, environmental, and historical processes in shaping genomic variation across the species’ range.

## 2. Materials and Methods

### 2.1 Field sampling and genetic data

Specimens of *Thrigmopoeus truculentus* were collected throughout the distribution range of the species. In total we collected 40 individuals from 11 locations across the WG throughout the extant distribution range. All specimens were preserved in ethanol and submitted to the Centre for Ecological Sciences museum, Bangalore, India. No collection permits were required for sampling, as *T. truculentus* is not scheduled under the Wildlife Protection Act, 1972 (latest amendment; Govt. of India, 2023), and all fieldwork was carried out outside protected areas.

Genomic DNA was extracted from leg tissues. Identity of specimens were confirmed according to keys provided in Siliwal & Molur, 2014 as well as through NCBI nucleotide-BLAST of a mitochondrial barcode region (details in the next section). We successfully generated genome wide SNPs data for 31 individuals from 11 locations, ensuring several individuals (2–4) were sequenced from every location. SNP data were generated through low-coverage whole genome sequencing at Eurofin Genomics (https://eurofinsgenomics.co.in/). Libraries were sequenced on a Novaseq6000 high-throughput sequencing platform (Illumina Inc.). On average, 3.5 GB of sequence data was generated per sample. Further details of the library preparation are provided in Supplementary Materials 1 (Section S1).

### 2.2 Mitochondrial phylogeny

A ∼750 bp fragment of the mitochondrial COI gene was amplified using the LCO1490/HCO2198 primer pair (Folmer et al., 1994) and sequenced using standard Sanger sequencing methods at https://www.barcodebiosciences.com/. Forward and reverse reads were inspected, trimmed, and aligned using MUSCLE algorithm (Edgar, 2004) in MEGA v10.0 (Kumar et al., 2018). All alignments were translated to amino acids to screen for indels and stop codons. A phylogeny was subsequently inferred using Bayesian inference in BEAST v2.5 (Bouckaert et al., 2019) using an uncorrelated relaxed molecular clock and a partition-specific substitution model, with MCMC chains run to convergence (10 million generations sampled every 1000 steps) and the maximum clade credibility tree was obtained.

### 2.3 Bioinformatics pipeline and data processing

Raw Illumina reads from all 31 individuals were subjected to a standardized quality-control and processing workflow prior to variant calling. Initial quality assessment of paired-end FASTQ files was performed using FastQC v0.11.9 (Brown et al., 2017) to inspect per-base quality trends, adapter contamination, and other sequencing artifacts. Adapter removal and quality trimming were conducted with Trimmomatic v0.39 (Bolger et al., 2014), employing a sliding-window approach (4 bp window, Q≥25) and retaining only reads ≥50 bp after trimming while discarding unpaired reads. Only paired, trimmed reads were retained for downstream analyses.

Filtered reads were aligned to the closest available tarantula reference genome, *Chaetopelma lymberakisi* (*GCA_964291345.1*), using BWA-MEM v0.7.17 (H. Li & Durbin, 2009). Alignment files were processed with SAMtools v1.17 (H. Li et al., 2009) for conversion, sorting, indexing, and duplicate removal. To avoid artificial inflation of allele counts in overlapping regions of paired reads, we further removed overlapping fragments using bamUtil clipOverlap v1.0 (Jun et al., 2015). Quality statistics including average genome-wide coverage were computed using Qualimap v2.3.1 (Okonechnikov et al., 2016). After alignment and filtering, samples retained a mean coverage of 0.08× (range: 0.1319–0.0213×; Supplementary Material 1, Table S1), suitable for genotype-likelihood–based inference under low-coverage conditions.

Genome-wide SNP calling was carried out using ANGSD v0.940 (Korneliussen et al., 2014), which calculates genotype likelihoods while accounting for sequencing uncertainty and low-coverage. Analyses were restricted to autosomal regions defined by the reference genome. SNPs were retained under specific filtering criteria to minimize false-positive variants: minimum mapping quality ≥20, base quality ≥20, exclusion of triallelic sites (-skipTriallelic 1), minor allele frequency ≥0.01 (-minMaf), presence in ≥80% of individuals (-minInd 24), and depth thresholds scaled to the sample number (-setMinDepth 31, -setMaxDepth 186). To remove potential sequencing and mapping artifacts, only uniquely mapped reads with properly paired alignments were retained (-uniqueOnly 1 - only_proper_pairs 1 -remove_bads 1). SNP discovery used a likelihood-ratio test with a conservative significance level (*p*<1e-6).

To minimize linkage disequilibrium (LD) bias for downstream population genomic analyses, we pruned linked variants using ngsLD v1.1.3 (Fox et al., 2019). Pairwise LD values were estimated from ANGSD-derived genotype likelihoods within 50 kb windows (--max_kb_dist 50), retaining only SNPs with low LD weight (--min_weight 0.2). A final LD-pruned SNP set was generated by filtering the BEAGLE-format genotype likelihood file, producing a dataset of 377483 high-quality, unlinked SNPs, which was used for all the downstream analyses.

### 2.4 Population structure analysis

We investigated population genetic structure using complementary genotype-likelihood-based approaches. First, we inferred major axes of genomic variation using PCAngsd v1.0 (Korneliussen et al., 2014), which estimates a pairwise covariance matrix directly from genotype likelihoods. Principal components were extracted in R (v4.1.2; R Core Team 2021) and visualized using *ggplot2* (Wickham, 2016), providing a model-free inference of clustering patterns among individuals.

Second, we estimated individual ancestry proportions using NGSadmix v3.2 (Skotte et al., 2013) across values of *K* = 1–6, with 15 independent replicates for each *K*. The maximum value of *K* was set to six as the mitochondrial phylogeny showed six broad clades (Results 3.1). Support for alternative clustering solutions was evaluated using both the Δ*K* method (Evanno et al., 2005) and the cross-validation (CV) error across replicates. Replicate results for the best-supported *K* were averaged and displayed as admixture barplots in *ggplot2*.

### 2.5 Calculation of genetic differentiation

To assess genetic differentiation among populations, we calculated pairwise *F_ST_* values between 11 populations using ANGSD v0.940 (Korneliussen et al., 2014) and realSFS (Nielsen et al., 2012). First, we generated a set of high-confidence shared SNPs across all individuals by estimating genotype likelihoods (-GL 1) and site allele frequencies (-doMaf 1), applying filters (SNP_pval ≤ 1e−6, minQ 20, minMapQ 20, uniqueOnly 1, only_proper_pairs 1, minInd 24, minMaf 0.01, and depth thresholds 31 ≤ depth ≤ 186). Coordinates of retained SNPs were extracted, deduplicated, and indexed to produce a shared-sites SNP panel used consistently across downstream analyses.

Population-wise BAM files were organized based on sampling localities (11), and site allele frequency likelihoods (-doSaf 1) were estimated separately for each population while restricting analyses to the shared SNP set (autosomal). Because no reliable outgroup genome is available for this species to confidently infer ancestral states, joint site frequency spectra (SFS) were folded to avoid incorrect assignment of derived alleles. Folded 2D-SFS were then generated for all population pairs. These 2D-SFS estimates were used to calculate pairwise *F_ST_* values. For comparative and spatial analyses, *F_ST_* values were also transformed into linearized *F_ST_* (*F_ST_* / (1 − *F_ST_*)). The final matrix of pairwise population differentiation was used to evaluate patterns of spatial genetic structure.

### 2.6 Assessment of genetic diversity

We quantified per-individual genetic diversity by estimating (i) the proportion of heterozygous sites per sample and (ii) individual inbreeding coefficients (F), using genotype likelihood–based methods.

To estimate heterozygosity, ANGSD v0.940 (Korneliussen et al. 2014) was used to identify a panel of shared SNPs across all individuals under the similar quality and depth filtering parameters in section 2.4. Genotype likelihoods were then calculated for each sample independently while restricting analysis to this shared SNP set. For each individual, we generated a folded 1-dimensional site frequency spectrum (1D-SFS) using realSFS (Nielsen et al., 2012), which does not require knowledge of the ancestral allele state. In a folded SFS for a single diploid individual, the middle frequency bin corresponds to heterozygous genotypes, whereas the two ends represent homozygous states. Because folding collapses symmetric allele configurations, the third bin was expected to be zero in this context and did not contribute to estimates. Individual heterozygosity (H_e_) was therefore computed as the proportion of sites assigned to the heterozygous bin relative to all informative sites retained for that sample.

We additionally estimated genome-wide inbreeding coefficients (F) using ngsF v1.2.0 (Vieira et al., 2013), which infers probabilistic deviations from Hardy-Weinberg equilibrium directly from genotype likelihoods. ANGSD was used to generate GLF likelihood files (-doGlf 3), and ngsF was run jointly across all samples using the same shared SNP panel. Per-individual F values were extracted from the resulting binary parameter file, where positive coefficients indicate elevated autozygosity consistent with inbreeding or reduced effective population size.

### 2.7 Ecological niche modelling

To evaluate habitat suitability and historical climatic stability across the distribution of *Thrigmopoeus truculentus*, we compiled occurrence records from our field surveys and supplemented them with additional verified records from GBIF (GBIF.org, 2025). All points were visually inspected for accuracy, and duplicate records were removed. Multiple records within a 1 km^2^ radius of each other were trimmed to ensure that each predictor raster cell contained no more than one presence record. After final filtering, 32 unique georeferenced presence points were retained for modelling (Table S2; Supplementary Material 1).

Environmental predictors were assembled from CHELSA climate datasets at 30 arc-second resolution, including present-day climate and paleoclimate reconstructions spanning 0–21 kya at ∼1000 yr intervals (Karger et al., 2023). All the raster layers were clipped to the extent of Western Ghats. Bioclimatic variables were extracted for all occurrence points, and highly correlated variables were removed using a Pearson correlation threshold of |r| ≥ 0.7. The final predictor set included eight variables: bio1, bio3, bio4, bio9, bio12, bio15, bio18 and bio19.

Species distribution models were implemented in R (R Core Team 2021) using the SDMtune package (Vignali et al., 2020) based on the Maxent maximum entropy algorithm (Phillips et al., 2006). A total of 10,000 background points were randomly sampled across the accessible area, and models were trained using an 70/30 train–test split. Model performance was assessed using area under the receiver operating characteristic curve (AUC), and jackknife tests were used to evaluate variable importance. The best-performing model was projected onto 20 paleoclimate layers representing conditions since the Last Glacial Maximum (LGM). The resulting 21 suitability timeseries surfaces were aggregated, and the Western Ghats layer was placed. The distribution range of the species was divided into 0.5 latitudinal bands (11 bands). For each band, temporal trend was summarized using mean, standard deviation, and coefficient of variation of suitable habitats as measures of regional habitat stability.

### 2.8 Predictors of genetic differentiation and genetic diversity

To identify the drivers of genetic differentiation among populations of *T. truculentus*, we modeled variation in pairwise genetic divergence using generalized linear mixed-effects models (GLMMs) implemented in R package lme4 (Bates et al., 2015). Pairwise linearized *F_ST_* was used as the response variable. Four biologically informed predictors were evaluated as fixed effects:

i. geographic distance — testing isolation-by-distance,
ii. difference in elevation — testing effects of topographic heterogeneity,
iii. difference in mean habitat suitability since the Last Glacial Maximum (LGM) — testing isolation-by-environment,
iv. difference in temporal variation of habitat suitability — testing effects of long-term climatic instability.

Because *F_ST_* is a pairwise measure with inherently non-independent observations, population identity for each pair member (Sample1 and Sample2) was included as a random effect. All continuous predictors were z-standardized prior to analysis to ensure comparability. We constructed five models with alternative predictor sets reflecting competing hypotheses, including a null model with random effects only (Table 1). Model fitting was performed using lme4, and model assumptions and residual structure were validated using DHARMa (Hartig F, 2024). Relative model support was evaluated using likelihood ratio tests against the null model and AIC scores using the package MuMIn (Bartoń K, 2025). Bioclimatic variables were not included directly because they are strongly collinear with the ENM-derived mean suitability estimates.

**Table 1:**
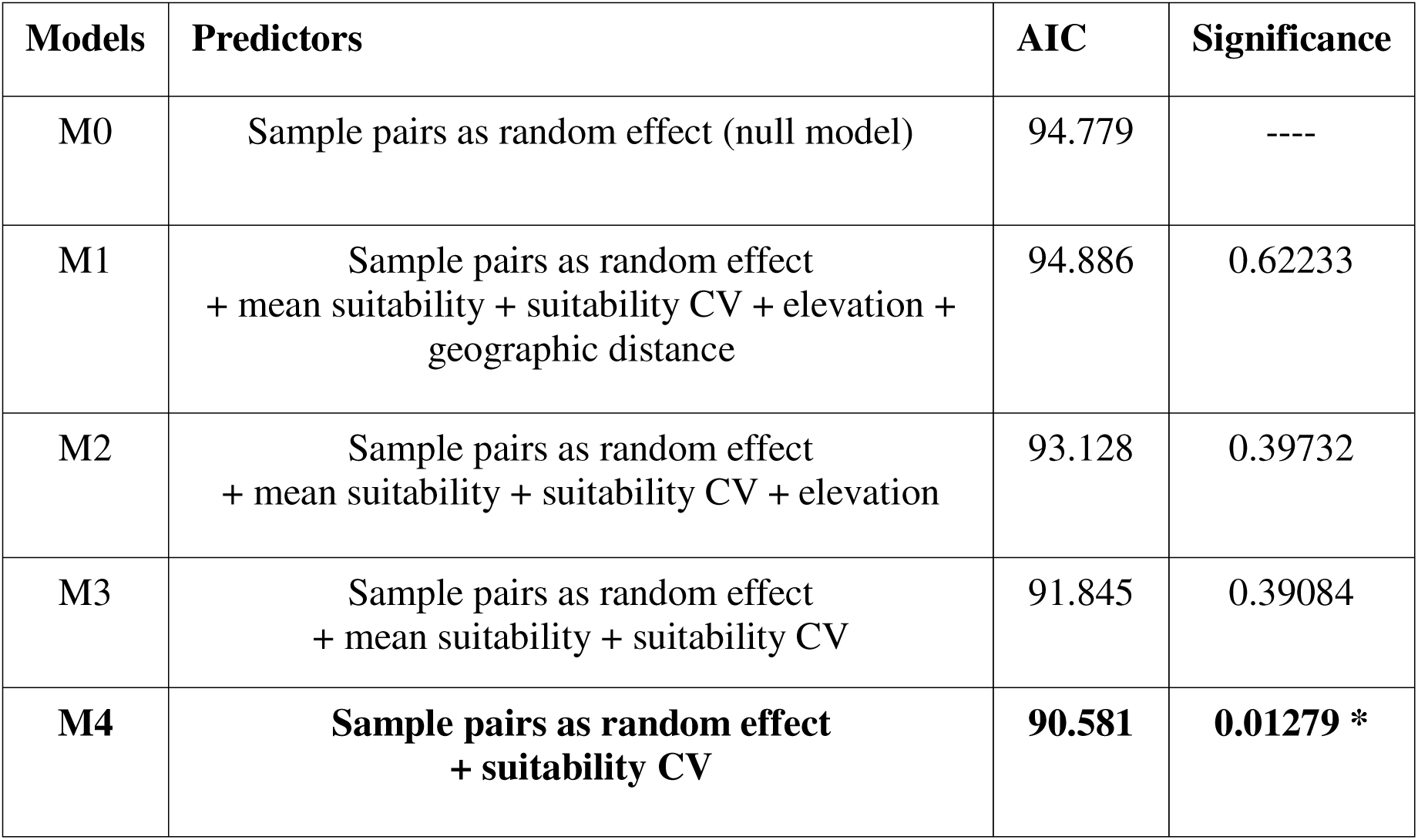
GLMM results with pairwise linearized *F_ST_* as response and mean suitability (isolation-by-environment), suitability CV (stability of habitat suitability), elevation (topographic heterogeneity) and geographic distance (isolation-by-distance) as predictors. The best model is highlighted in bold.

For per-individual genetic diversity, we used two response variables (i) the proportion of heterozygous sites and (ii) individual inbreeding coefficients (F). We examined a predictor set including mean habitat suitability, temporal variation in habitat suitability, elevation, and selected bioclimatic parameters from CHELSA. To account for multicollinearity among climatic predictors and improve variable selection, we implemented a LASSO regression framework using glmnet (Friedman et al., 2010) package, with coefficients estimated at the λ minimizing cross-validation error. Predictors retained by LASSO were subsequently evaluated in ordinary least squares regression models to quantify effect sizes and variance explained (adjusted R²). Model predictions and uncertainty were visualized using linear fits with 95% confidence intervals.

### 2.9 Inferring demographic history

To reconstruct historical demographic trajectories, we estimated changes in effective population size (Ne) over time for the species as a whole and for three groups (north, central, south) separately. Groups were defined based on a combination of ancestry proportions inferred from NGSadmix analyses, sampling locations, and ecological niche modeling, which identified three broad regions with distinct paleoclimate dynamics. Each group encompassed four populations.

We first generated folded site frequency spectra (SFS) for each dataset using ANGSD v0.935 and realSFS (Nielsen et al., 2012). The species-wide analysis included 31 individuals, whereas sub-population analyses included 10 or 11 individuals per region (north, central, south). A common set of genomic sites were retained as per the filtering conditions mentioned in section 2.4.

These SFS were then used as input for Stairway Plot v2 (Liu & Fu, 2020). For all analyses, we specified a mutation rate of 5 × 10 per site per generation based on estimates for other spiders (Escuer et al., 2024; Ma et al., 2025) and a generation time of 4 years (Schwerdt et al., 2021), based on literature. To improve robustness, each analysis included 200 bootstrap replicates and model parameters tuned to the number of sequences in each dataset.

### 2.10 Sensitivity analysis: Effects of variable sequencing effort

Differences in sequencing depth among individuals can introduce bias in genotype-likelihood–based population genomic analyses, particularly when working with low-coverage whole-genome data. In our dataset, sequencing coverage varied by approximately six-fold across samples (0.02×–0.13×). To evaluate whether this variation influenced population structure and genetic diversity estimates, we performed a downsampling analysis.

All higher-coverage BAM files were randomly downsampled to match the lowest-coverage sample using SAMtools (samtools view -s), ensuring comparable read depth across individuals. Genotype likelihood estimation and SNP calling were repeated in ANGSD using the same filtering parameters described in Section 2.2, resulting in a reduced SNP panel (318851 SNPs). We then re-ran the PCAngsd, NGSadmix, and genetic diversity analyses (heterozygosity and inbreeding coefficients) on the downsampled dataset.

Results from the downsampled analyses were qualitatively compared to those generated from the full dataset to determine whether variable sequencing coverage impacted the main biological conclusions.

## 3. Results

### 3.1 Population structure

Mitochondrial phylogeny recovered six broad geographic clades with strong posterior support (PP = 1; Fig. 2a). Individuals from Amboli (northernmost locality) formed a distinct clade, while samples spanning Castlerock/Bhimgad and Sirsi grouped into a second clade. A third clade included Coorg, Chikmagalur and Sakleshpur populations. The fourth clade included Agumbe and Keemangundi populations. Mananthawady and Honey valley populations formed the fifth clade, while the Wayanad specimens fell in the sixth clade. Despite strong support for terminal clades, relationship between these clades showed low support (PP = 0.34–0.59), indicating uncertainty in deeper mitochondrial relationships.

**Figure 2:**
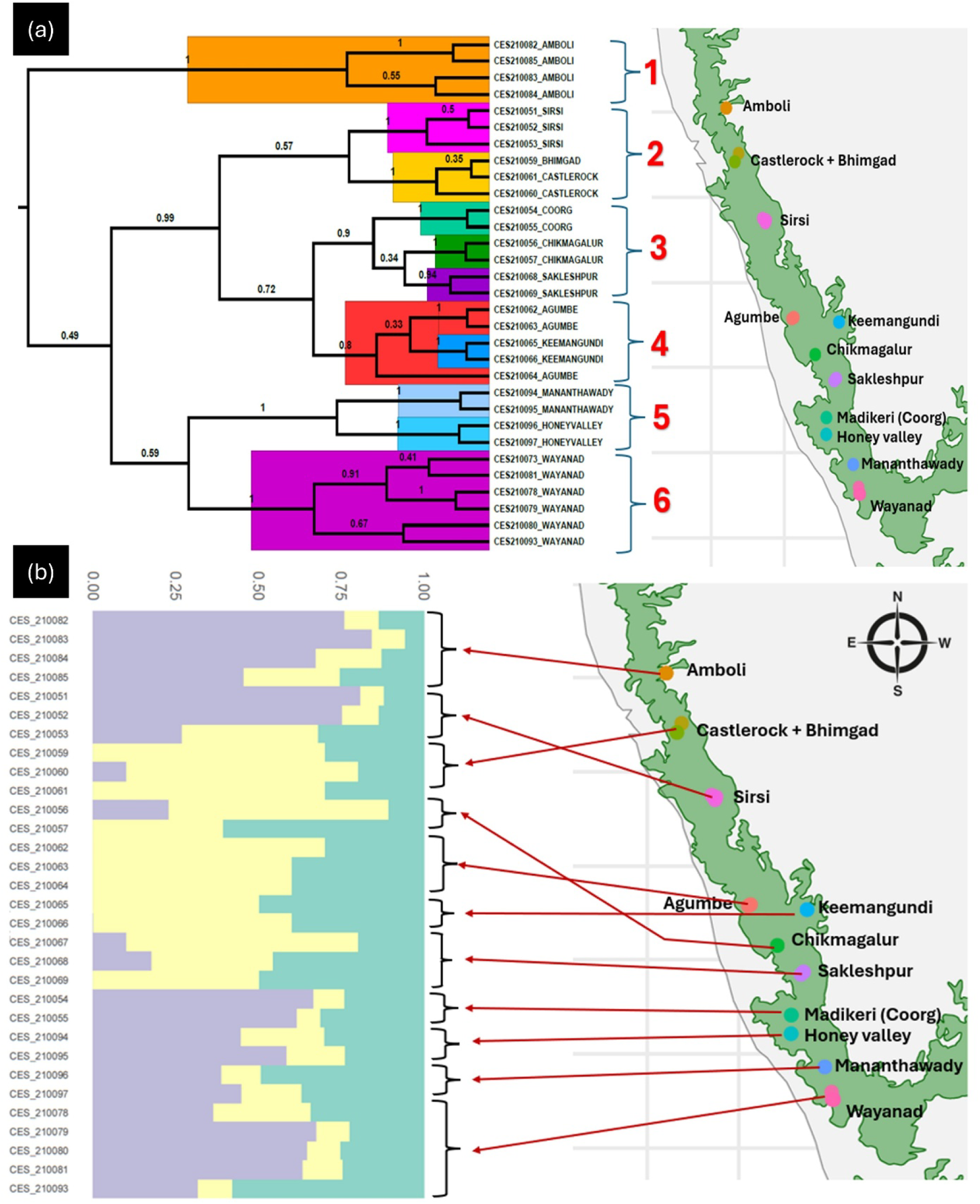
a) The mitochondrial phylogeny and b) the admixture plot for k = 3 along with the sampling locations.

In contrast, nuclear SNP data revealed only weak genetic structure. Both ΔK and cross-validation error identified K = 3 as the best-supported model in NGSadmix (Fig. S1, Supplementary Material 2). However, ancestry coefficients showed high admixture across all individuals, with no population exhibiting a clearly discrete gene pool (Fig. 2b). A subtle pattern emerged: central Western Ghats populations (Keemangundi, Agumbe, Chikmagalur, Sakleshpur) displayed relatively higher percentage of yellow ancestry, whereas northern (Amboli, Sirsi) and southern (Madikeri, Mananthawady, Honey Valley, Wayanad) populations showed higher proportions of purple ancestry (Fig 2b). Some northern individuals (Castlerock, Bhimgad) appeared similar to the central ones, suggesting genetic continuity across the range. Admixture plots for K = 2 and 4 displayed broadly a similar pattern as K = 3 (Figure S2 and S3; Supplementary Material 2),

PCAngsd findings were consistent with the admixture results. PC1 (9.04% variance) separated the central samples from a cluster of northern + southern individuals, while PC2 explained little structure (3.58%) but hinted at mild north–south differentiation (Fig. 3). Considerable overlap among clusters again indicated no strong population boundaries.

**Figure 3:**
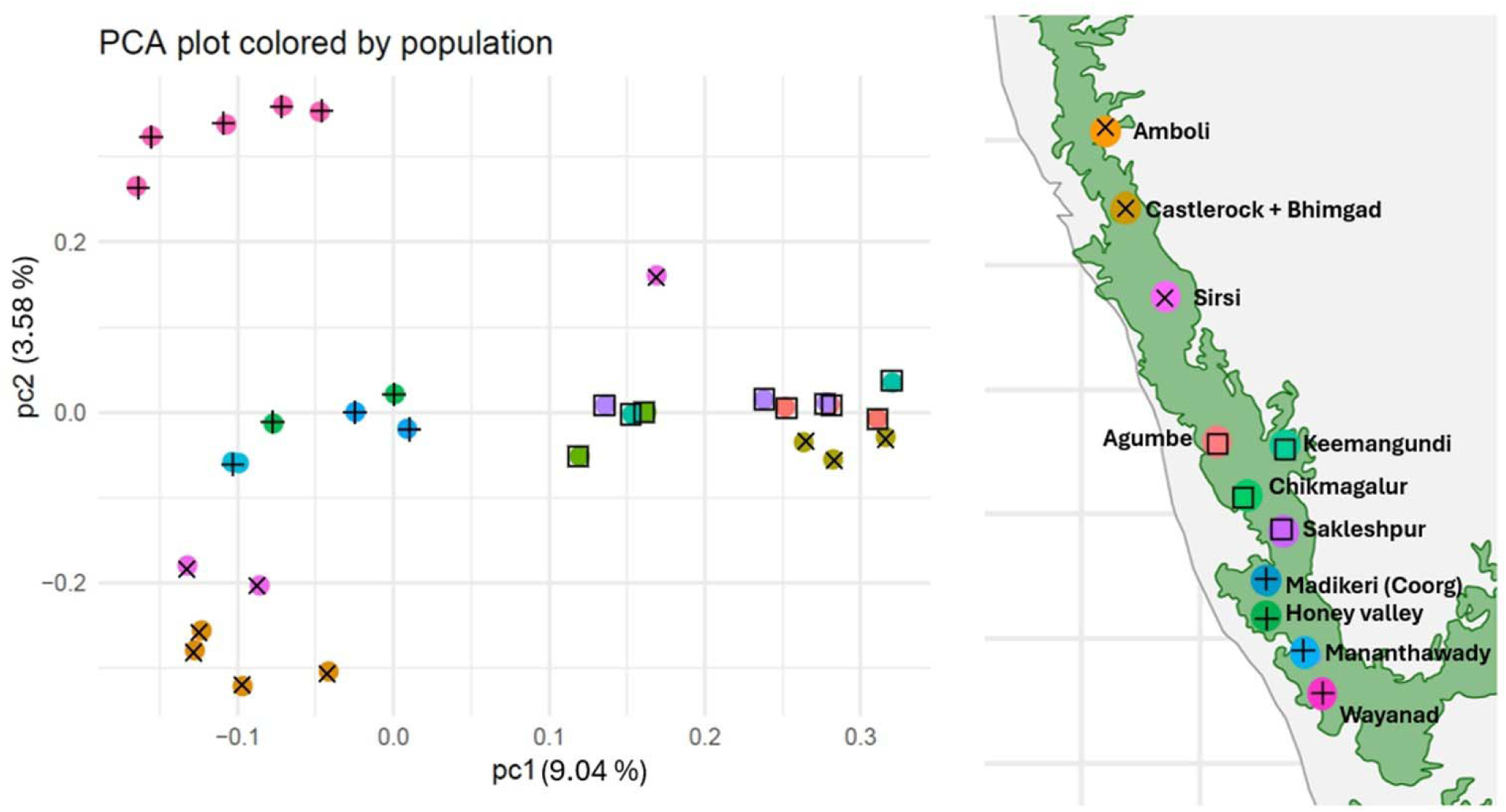
PCA plot along with the proportion of variation explained by each axis. Samples are marked according to their location. Crosses indicate samples from the northern region, plus indicate samples from the southern part, and boxes denote samples from the central region.

Taken together, mitochondrial and nuclear markers portray differing patterns: mtDNA suggests a latitudinal phylogeographic signal, whereas genome-wide SNPs reveal extensive gene flow and only weak, diffuse population structure. These patterns support predominantly continuous connectivity, with a modest signature of differentiation in the central part of the distribution.

### 3.2 Patterns of genetic differentiation and diversity

Pairwise *F_ST_* estimates indicated overall low levels of genetic differentiation among the sampled populations, with an average *F_ST_* = 0.029. Linearized FST values ranged from 0.000189 to 0.1476. A heatmap matrix of pairwise *F_ST_* values is presented in Figure S4 (Supplementary Material 2). The highest *F_ST_*estimates occurred in comparisons involving populations from the central region versus those from the northern and southern regions, reflecting a degree of genetic distinctiveness in some of the central populations. In contrast, pairwise *F_ST_* values among northern and southern populations remained consistently low. It is important to emphasize, however, that the overall differentiation is minimal.

Line plots of heterozygosity and inbreeding coefficients are shown in Figure S5 (Supplementary Material 2). Patterns of genetic diversity revealed comparatively high heterozygosity across most locations, except within the central populations (Agumbe, Keemangundi, Chikmagalur and Sakleshpur), which exhibited slightly lower heterozygosity. The inbreeding coefficient (F) showed the opposite trend: although inbreeding levels remained low overall, modest increase was observed in the central region (Agumbe, Keemangundi, Chikmagalur and Sakleshpur), peaking around F ≈ 0.015, which is higher relative to the other populations.

Together, these results suggest that while genetic diversity is generally high and inbreeding is low across the species’ range, populations from the central region show relatively reduced diversity and increased inbreeding, indicating potential genetic structuring and different demographic patterns relative to the northern and southern populations.

### 3.3 Paleohabitat dynamics

The climatic niche model for *Thrigmopoeus truculentus* showed high predictive accuracy, with an AUC of 0.978. Jackknife analysis identified Bio15 (precipitation seasonality), Bio4 (temperature seasonality), and Bio1 (annual mean temperature) as the most influential predictors of the species’ ecological niche.

Time-series habitat suitability projections from the present to the Last Glacial Maximum (LGM) revealed spatially heterogeneous paleohabitat patterns across the species’ range (Figure 4a). The northern and southern regions of the species’ range maintained consistent habitat suitability through time. In contrast, the central region—encompassing Agumbe, Keemangundi, Sakleshpur, and Chikmagalur, displayed marked temporal fluctuations in suitability. For example, periods such as ∼2 ka, ∼9 ka, and ∼13 ka presented highly suitable conditions in the central region, whereas at other times (e.g., present day, ∼4 ka, and ∼11 ka) suitability dropped sharply (Figure 4a).

**Figure 4:**
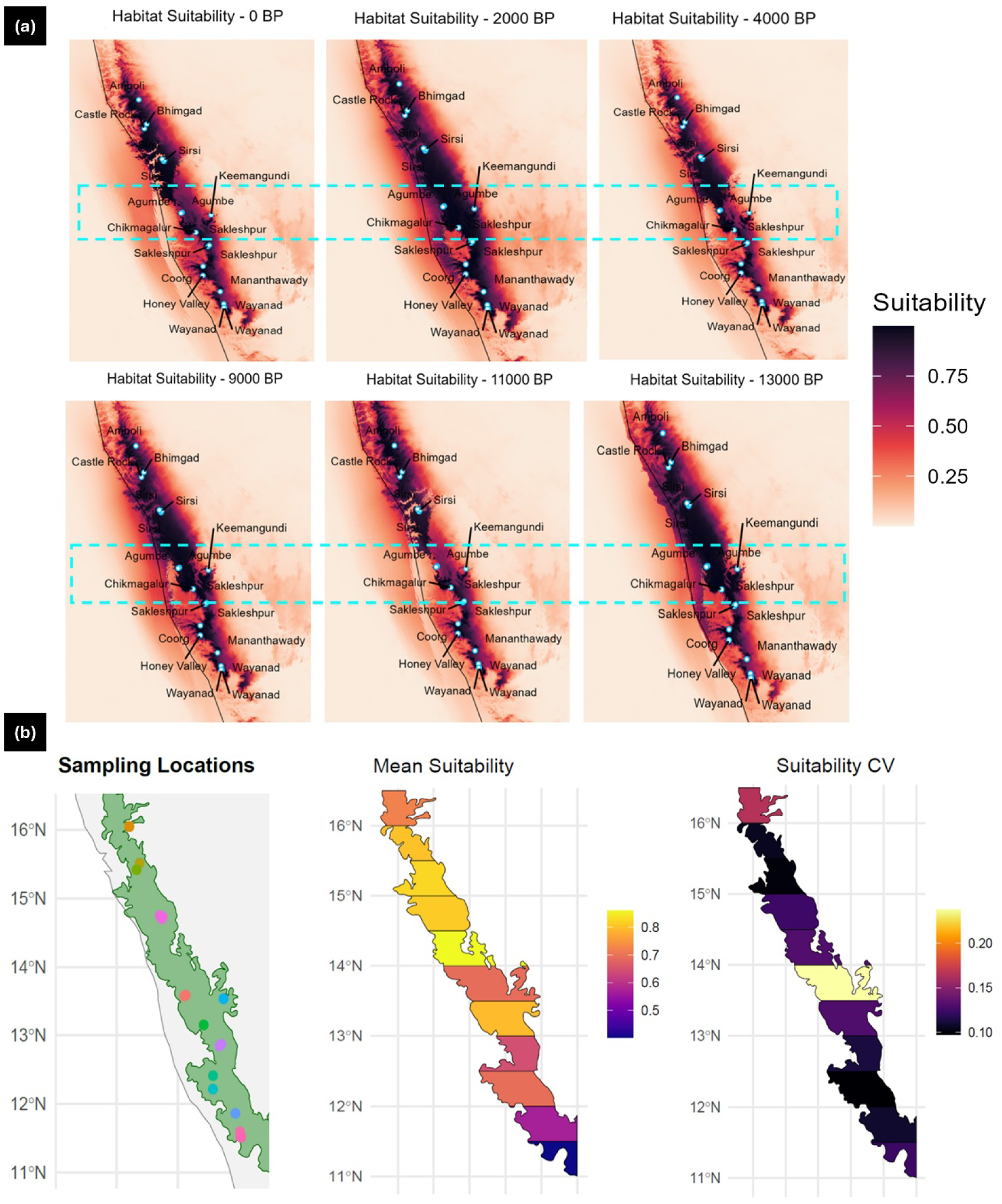
a) Habitat suitability rasters for some selected time points. The region of fluctuating suitability is highlighted with a rectangle and b) Paleohabitat suitability statistics presented for every 0.5□ latitude bands across the distribution range of *T. truculentus* (sampling locations, mean suitability per band since LGM, and coefficient of variation of suitability since LGM).

To quantify these dynamics, we extracted key temporal habitat stability metrics across 0.5 latitudinal bands within the Western Ghats boundary (since the species is endemic to the region). As shown in Figure 4b, both the standard deviation and coefficient of variation of habitat suitability were substantially higher between 13–14°N, corresponding to the central populations. Meanwhile, latitudes spanning the northern (≈14–16°N) and southern (≈11–13°N) parts of the range exhibited low variability, indicating long-term habitat stability.

This pattern suggests that the central part of the distribution range experienced episodic environmental instability, likely exposing populations to periodic demographic stress, contraction events, or fluctuating population sizes over late-Quaternary climatic cycles. Such instability can leave stronger genetic footprints than more stable environments, providing a plausible explanation for why central populations exhibit reduced heterozygosity, slightly higher inbreeding, and distinct genomic signatures, despite the overall low genetic differentiation across the species.

All paleohabitat suitability maps for the full time series are provided in Supplementary Material 3.

### 3.4 Predictors of genetic differentiation and genetic diversity

Generalized linear mixed-effects models revealed that historical stability of suitable habitat, quantified as the coefficient of variation (CV) in habitat suitability since the LGM, was the strongest and only significant predictor of genetic differentiation among populations (Table 1). Model comparison indicated that the model including CV of suitability (Model 4) provided a significantly better fit than the null expectation, explaining ∼85% of the variation in pairwise *F_ST_*values. Geographic distance, elevation, and mean suitability across time did not meaningfully improve model performance. CV of suitability showed a positive association with pairwise differentiation (β = 0.226, SE = 0.084), demonstrating that populations occurring in areas where suitable habitat fluctuated strongly through time are more genetically differentiated, whereas populations in consistently suitable regions are genetically similar (Fig. 5a). Diagnostic residual checks confirmed excellent model performance (Figure S6, Supplementary Material 2), and effect-size estimates from the full model (M1) reinforced CV of suitability as the only impactful predictor (Fig. 5b).

**Figure 5:**
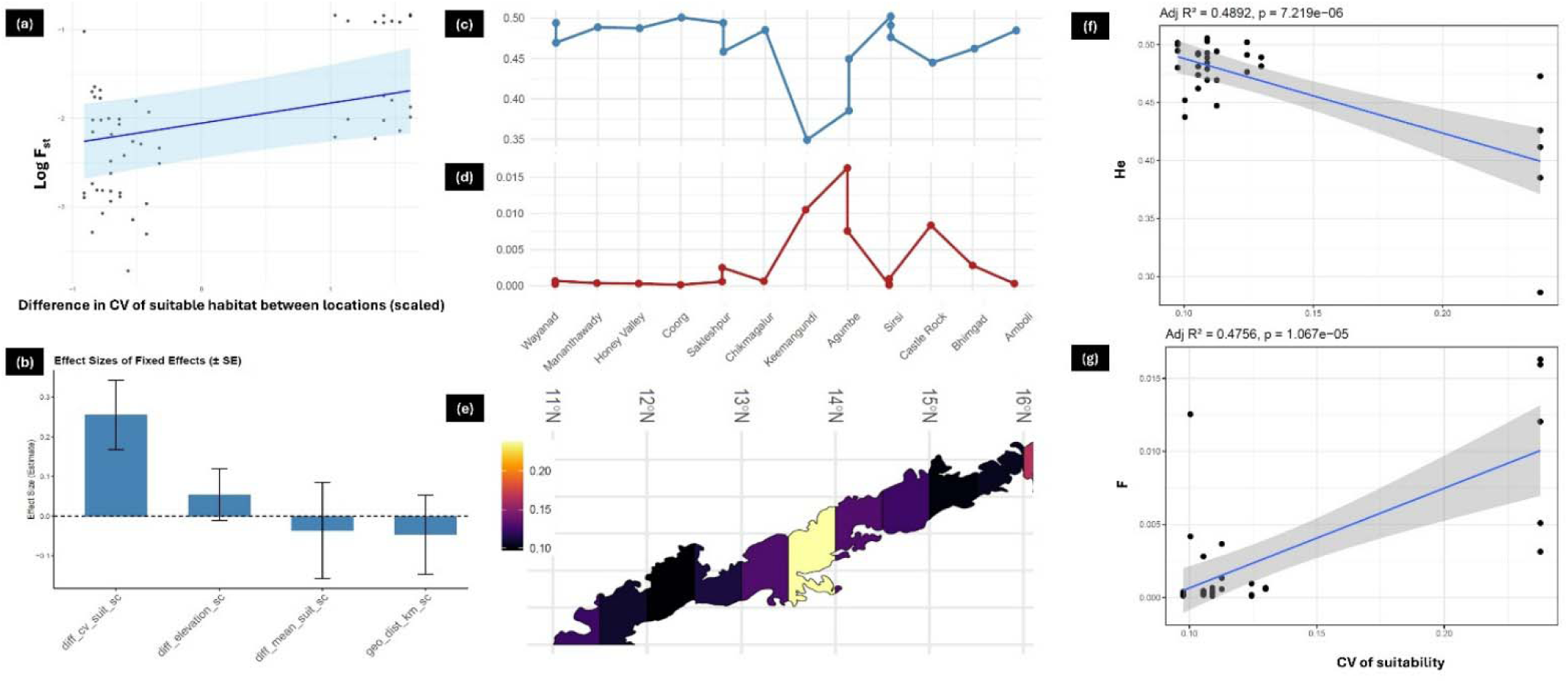
a) The relationship between genetic differentiation and difference in CV of suitability across sampling locations b) Estimated effect sizes of four predictors according to the most inclusive GLMM model c) and d) show the trend of heterozygosity and inbreeding coefficient values respectively across locations and compared against the latitude CV suitability bands (e). The ordinary linear regression relationship between CV of suitability and heterozygosity as well as inbreeding coefficients are shown in panel (f) and (g) respectively.

Patterns of genetic diversity showed a similar signal. LASSO regression identified CV of suitability as the only retained predictor for both the proportion of heterozygous sites and inbreeding coefficients across individuals. Populations located around 13–14°N, where suitable habitat has been least stable over time, exhibited the lowest heterozygosity and highest inbreeding (Figs. 5c–e). Linear regression models further supported these relationships: CV of suitability had a strong negative effect on heterozygosity (R²adj = 0.49; Fig. 5f) and a strong positive effect on inbreeding (R²adj = 0.47; Fig. 5g).

Together, these analyses show that temporal instability of suitable habitat, rather than simple geographic distance or topographic variation, is the principal factor shaping both genetic differentiation and diversity in *T. truculentus* across the Western Ghats.

### 3.5 Demographic history

Demographic reconstruction for the full dataset revealed a gradual decline in effective population size (Ne) throughout the Pleistocene, followed by marked stabilization around the Last Glacial Maximum (Fig. S7, Supplementary Material 2). Since the LGM, *Thrigmopoeus truculentus* has maintained a relatively stable Ne, with a median estimate of ∼8×10 individuals.

Group-wise analyses further uncovered region-specific demographic histories (Fig. 6). When compared alongside paleohabitat dynamics and admixture patterns (Fig. 6a–c), the northern lineage exhibited a trajectory similar to the species-wide pattern, maintaining stable population sizes since the LGM (median Ne ∼2.6×10). The southern lineage also remained broadly stable, showing only a mild post-LGM decline followed by stability ∼10 kya (median Ne ∼2.9×10).

**Figure 6:**
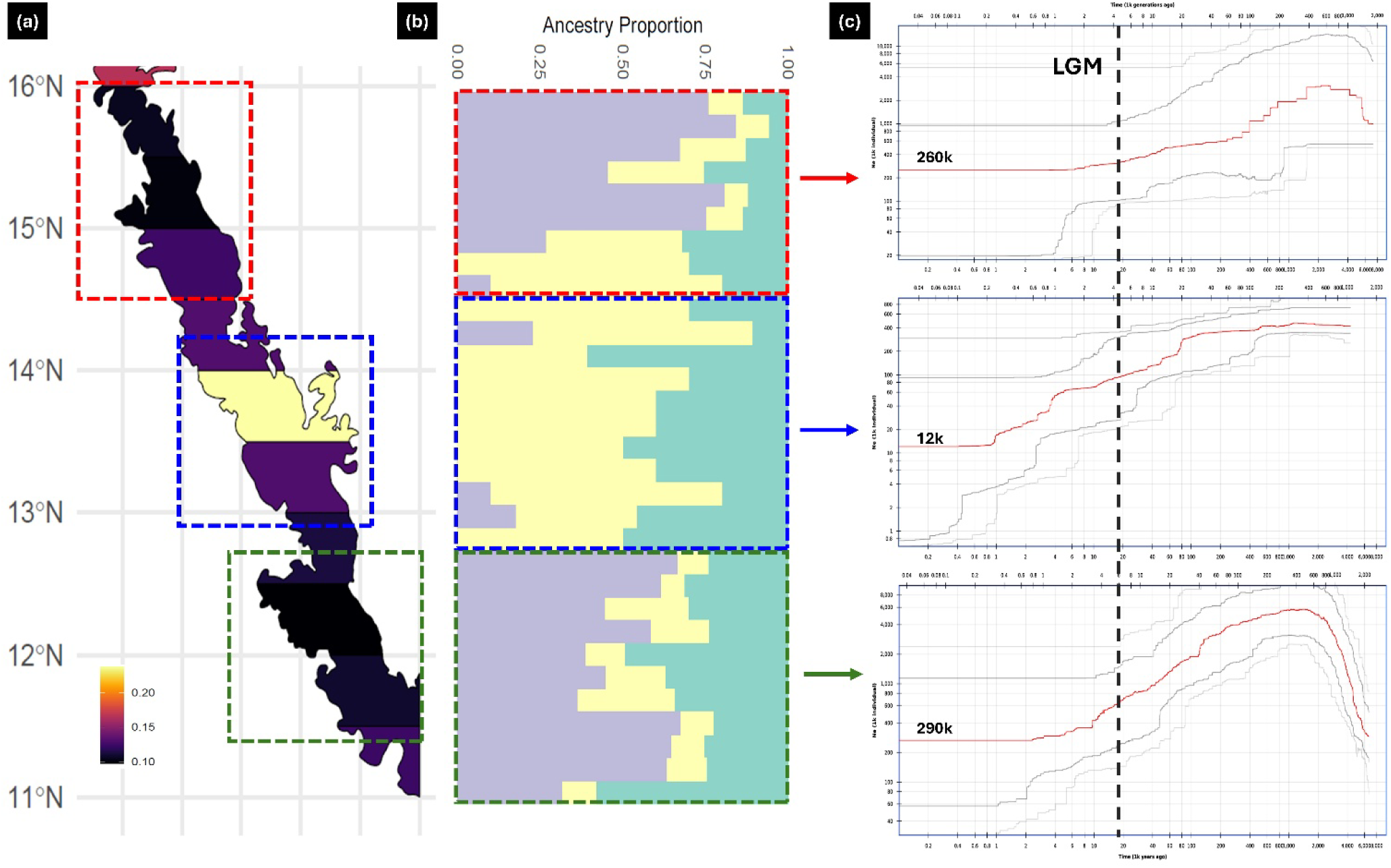
The stability of suitable habitats (a), admixture proportions (b) and the effective population size trajectories (c) are compared side by side for three groups. Estimated extant effective population sizes are mentioned in panel (c).

In contrast, the central lineage displayed a pronounced and prolonged population decline, beginning ∼100 kya and continuing well beyond the LGM until ∼1 kya. This group also maintained substantially lower population sizes overall (median Ne ∼1.2×10), consistent with reduced genetic diversity and elevated inbreeding observed in previous analyses.

Collectively, these demographic reconstructions highlight a long-term demographic stability in the northern and southern regions, likely reflecting persistent habitat suitability, whereas the central lineage experienced sustained demographic instability, in line with stronger fluctuations in habitat suitability and associated signatures of genetic erosion.

### 3.6 Sensitivity analysis

Population structure and diversity patterns inferred from the downsampled dataset were highly consistent with the main analyses. PCAngsd results again showed weak structure, with PC1 (9%) separating central populations from the northern and southern groups, mirroring the primary dataset (Figure S8; Supplementary Material 2). Similarly, NGSadmix identified K = 3 as the optimal clustering level. The ancestry profiles remained highly admixed, with the same broad pattern of genetic similarity between northern and southern samples and subtle distinctiveness of central populations (Figure S9; Supplementary Material 2).

Genetic diversity metrics were also robust to the depth-control procedure. Although heterozygosity estimates were high overall (expected due to reduced SNP filtering under lower depth), central populations still exhibited comparatively lower heterozygosity. Inbreeding coefficients likewise showed the same latitudinal trend observed previously, with elevated inbreeding in central populations relative to northern and southern regions (Figure S10; Supplementary Material 2).

Together, these results indicate that the spatial patterns of genetic structure and diversity recovered in this study are not artifacts of unequal sequencing depth but reflect true underlying biological signal.

## 4. Discussion

This study provides the first population genomic assessment of a Western Ghats–endemic species, delivering novel insights into evolutionary processes in a globally significant biodiversity hotspot. By integrating genome-wide SNPs with reconstructions of historical habitat suitability and demographic change, we reveal a unique case in which subtle but meaningful population structure has emerged despite the absence of major geographic barriers and the potential for panmixia. Our findings demonstrate that long-term fluctuations in the extent and stability of suitable habitats, rather than simple spatial isolation — can drive genomic differentiation and shape demographic resilience. This work illustrates how paleoclimatic processes can produce unexpected and cryptic population structure, emphasizing the broader relevance of these processes beyond a single regional taxon.

### 4.1 Temporal habitat stability as the primary driver of genomic differentiation

Population genomic analyses, including PCA and admixture inference, revealed generally weak structuring within *T. truculentus*, probably because of historical gene flow across its range. Yet, a particularly striking pattern emerged. Individuals from the northern and southern edges of the species’ distribution are genetically more similar to each other than either group is to individuals from the center of the range. The lack of a clear pattern following geographic proximity indicates that contemporary spatial separation alone cannot account for this structure. Importantly, the pattern remained even after accounting for possible technical bias, supporting a genuine biological signal.

To understand this pattern, we examined paleoclimate-informed habitat suitability across time. Habitat models extending from the Last Glacial Maximum to the present show that the central portion of the distribution experienced strong temporal fluctuations in habitat suitability. In contrast, both the northern and southern regions exhibited prolonged stability in suitable habitat conditions. Statistical analyses identified habitat stability, measured through the coefficient of variation in suitability over time, as the most influential predictor of genetic differentiation and diversity. Demographic reconstructions further support this interpretation, showing that central populations have experienced a prolonged decline in effective population size, whereas northern and southern populations stabilized following the LGM.

These results collectively point to a scenario in which *T. truculentus* likely maintained well-connected and largely panmictic populations throughout its historical distribution. However, repeated reductions in habitat suitability within the central region over the past several thousand years would have periodically decreased local population sizes. Such demographic instability likely caused reduced genetic diversity, increased inbreeding, and ultimately greater genetic differentiation in these central populations. The genomic signature observed today therefore reflects the cumulative impacts of fluctuating habitat suitability rather than isolation driven by distance or physical barriers. In contrast, the consistently suitable habitats to the north and south appear to have preserved genomic similarity between distant populations.

Our findings align with growing evidence that paleoclimatic changes have played a major evolutionary role in Western Ghats. Chaitanya et al. (2025) demonstrated that climatic transitions around the Coorg–Wayanad region can promote speciation in endemic reptiles even when geographic barriers are absent. Likewise, Bose et al. (2016) found the 13 to 14 degree latitude belt as a region of long-term climatic fluctuations (Fig 1), particularly concerning precipitation seasonality, a variable that also emerged among the strongest predictors of *T. truculentus* habitat suitability. These parallels suggest that climatic instability in the mid-latitude Western Ghats might act as a persistent ecological filter, shaping population genetic structure across multiple taxa.

Although climate-driven genetic structuring has been documented in other regions globally (de Pous et al., 2016; Fonseca et al., 2023; Lima et al., 2017; Morgan et al., 2011; Nogués-Bravo et al., 2010; Pearce-Higgins et al., 2015; Willis et al., 2004), the specific pattern uncovered here appears novel. The emergence of genetic divergence in the central region while the terminal populations remain genetically similar challenges the dominant view that iconic physical barriers such as the Palghat and Shencottah gaps are the primary generators of population structure in the Western Ghats. Instead, our results highlight the potential for dynamic paleoclimatic processes to create less obvious but equally influential evolutionary scenarios.

Taken together, this study introduces a new biogeographic hypothesis for the Western Ghats. The spatially variable climatic stability may produce more intricate patterns of isolation and connectivity than those implied by geography alone. Future population genomic studies across co-distributed species will be essential for testing whether such processes are widespread and for clarifying the extent to which long-term habitat stability has shaped the genetic diversity of taxa from this region.

### 4.2 Unique genetic structure relative to other mygalomorphs

Mygalomorph spiders, including tarantulas, trap-door spiders, and purse-web spiders, are usually characterized as sedentary arthropods with limited dispersal potential (Biswas et al., 2023; Foley et al., 2021; Monjaraz-Ruedas et al., 2024). Numerous phylogeographic studies across continents have supported this view, consistently reporting strong population genetic structure driven by restricted gene flow (Arnedo & Ferrández, 2007; J E Bond et al., 2001; Opatova et al., 2016; Opatova & Arnedo, 2014), often inferred largely from mitochondrial markers or a small number of nuclear loci. This broad pattern of high spatial genetic subdivision has been informally termed the “mygalomorph rule” (Monjaraz-Ruedas et al., 2024). However, recent genomic studies have shown that exceptions can occur, particularly in species capable of occasional long-distance dispersal or those that have experienced recent demographic shifts that prevent mutation–drift equilibrium (Hamilton et al., 2011; Monjaraz- et al., 2024).

Contrary to the expectations of the mygalomorph rule, *T. truculentus* exhibits strikingly low levels of population differentiation, with low *F_ST_*estimates and high admixture among populations. These results indicate substantial historical connectivity across its distribution. Despite ecological specialization, the continuous distribution of forested habitat in this region may have facilitated persistent gene flow (Reddy et al., 2016), maintaining genetic cohesion across populations. Local abundance and high effective population sizes may further buffer this species from genetic drift, reducing opportunities for differentiation to accumulate.

We also detected notable mito-nuclear discordance, which is best explained by male-biased dispersal. Females remain burrow-bound and sedentary for life, whereas males become wide-ranging during the mating season and can move over considerable distances, a behavior documented in multiple tarantula species (Ferretti et al., 2014; Janowski-Bell & Horner, 1999). Because mitochondria are maternally inherited, restricted female movement leads to stronger mitochondrial structure even when nuclear genomes remain mixed (Ellegren & Galtier, 2016; Vachon et al., 2018).

Taken together, *T. truculentus* represents an informative exception to widely assumed population genetic trends in mygalomorph spiders. However, it’s worth noting that *T. truculentus* does confirm the mygalomorph rule with respect to the patterns of mitochondrial genetic divergence. Importantly, such insights remain limited by a globally sparse dataset. As only a handful of mygalomorphs have been studied using genomic data, expanding the taxonomic coverage will be critical to assess whether exceptions like *T. truculentus* are rare anomalies or simply reflect our still-limited understanding of population processes in this group.

Further, our results align with recent comparative analyses showing that larger-bodied mygalomorphs tend to exhibit higher genetic diversity (Monjaraz-Ruedas et al., 2024). *T. truculentus*, a large tarantula species (4–5 cm total body length), shows high overall heterozygosity and relatively low inbreeding.

### 4.3 Conservation and taxonomic implications

*Thrigmopoeus truculentus* is currently listed as “Near Threatened” on the IUCN Red List, largely due to habitat loss and degradation associated with rapid developmental activities in the Western Ghats (Molur, 2015). Despite these concerns, many aspects of its ecology, including dispersal capability, local population density, and demographic stability in the wild — have remained poorly understood. Our study provides the first genome-wide assessment of population status in this endemic tarantula and reveals encouraging signals for its long-term persistence. We find that the species maintains a historically stable and relatively large effective population size since the Last Glacial Maximum, alongside high genetic diversity and low levels of inbreeding across most of its range. Only populations in the central part of the range (∼13–14 N) show evidence of reduced effective population sizes and stronger demographic fluctuations, making this region a potential focus for targeted monitoring.

These results suggest that *T. truculentus* remains broadly resilient, likely supported by relatively contiguous forest cover across its distribution, which has helped preserve connectivity and gene flow (REDDY et al., 2016). However, this should not be interpreted as diminishing existing conservation concerns. The species continues to play an important role in Western Ghats ecosystems as a dominant fossorial predator and ecosystem engineer (Formanowicz Jr. & Ducey, 1991; Siliwal & Molur, 2009). Ongoing land-use changes still pose a substantial risk, and efforts to preserve habitat continuity remain essential to prevent future erosion of genetic health.

Our findings also hold important taxonomic implications. Our mitochondrial phylogeny revealed multiple well-supported clades with notable sequence divergence, a pattern if interpreted in isolation, might suggest the presence of cryptic lineages within *T. truculentus*. However, mitochondrial markers are known to exhibit elevated lineage sorting and can strongly overestimate population divergence due to their haploid, maternally inherited, and non-recombining nature (Ellegren & Galtier, 2016; Monjaraz-Ruedas et al., 2024; Vachon et al., 2018). In addition to the pitfalls of mtDNA, traditional morphology-based taxonomic decisions in mygalomorphs can be misleading because phenotypic characters often evolve slowly, show high intraspecific variation, and lack clear diagnostic resolution (Ayoub et al., 2007; Jason E Bond et al., 2012; Lüddecke et al., 2018).

Genome-wide SNP data provides a contrasting and more comprehensive picture. Despite strong mitochondrial structuring, nuclear analyses demonstrate high admixture, extensive gene flow, and no evidence of reproductive isolation across the species’ range. These results strongly support the interpretation of *T. truculentus* as a single, genetically cohesive species with only subtle regional structure. Given these findings, we recommend caution regarding any proposed taxonomic subdivision of *T. truculentus* based solely on mitochondrial sequences or limited morphological traits. Higher-resolution nuclear genomic approaches should remain the standard for diagnosing independently evolving lineages in this group.

### 4.4 Limitations, uncertainties, and future research directions

While this study provides the first genome-wide insights into the evolutionary history of *T. truculentus*, several limitations should be acknowledged. First, because no reference genome currently exists for this species or its close relatives in the Western Ghats, we aligned reads to the most closely available tarantula genome. Although our downstream filtering and genotype-likelihood framework are specifically designed to handle uncertainty, future analyses based on a species-specific genome would improve mapping efficiency, SNP recovery, and spatial resolution of population structure.

Second, demographic reconstructions relied on low-coverage data and the site frequency spectrum, which inherently carry uncertainty. Although the median trajectories of the three regional groups differ significantly, particularly the pronounced decline in the central populations. However, overlap in the confidence intervals indicates that these patterns should be interpreted with caution and validated using deeper sequencing or complementary demographic inference methods.

Despite these uncertainties, our core findings, i.e. high overall connectivity, localized genomic erosion in the central Western Ghats, and strong links between habitat stability and genomic variation—are strongly supported across multiple independent analytical frameworks. Moving forward, generating a high-quality reference genome, and conducting comparative genomic studies across additional Western Ghats endemics will be valuable steps toward assessing whether the processes uncovered here represent a broader regional pattern.

## Data availability statement

All the raw sequencing reads were submitted to the National Center for Biotechnology Information (NCBI) under the BioProject number PRJNA1389812.

