## Supplementary material 2 for "Spatiotemporal variation in habitat suitability predicts genomic diversity and structure in a Western Ghats endemic Tarantula (*Thrigmopoeus truculentus*)"

**Figure S1:** CV error and ΔK plots. K=3 and K=5 shows almost comparable CV error trend but K=3 outperforms K=5 in terms of ΔK.

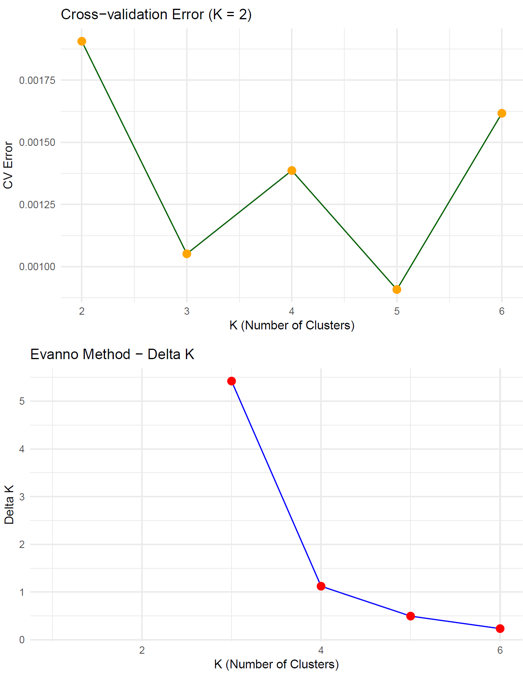

**Figure S2:** Admixture plot for K=2. Samples are plotted from north to south (left to right).

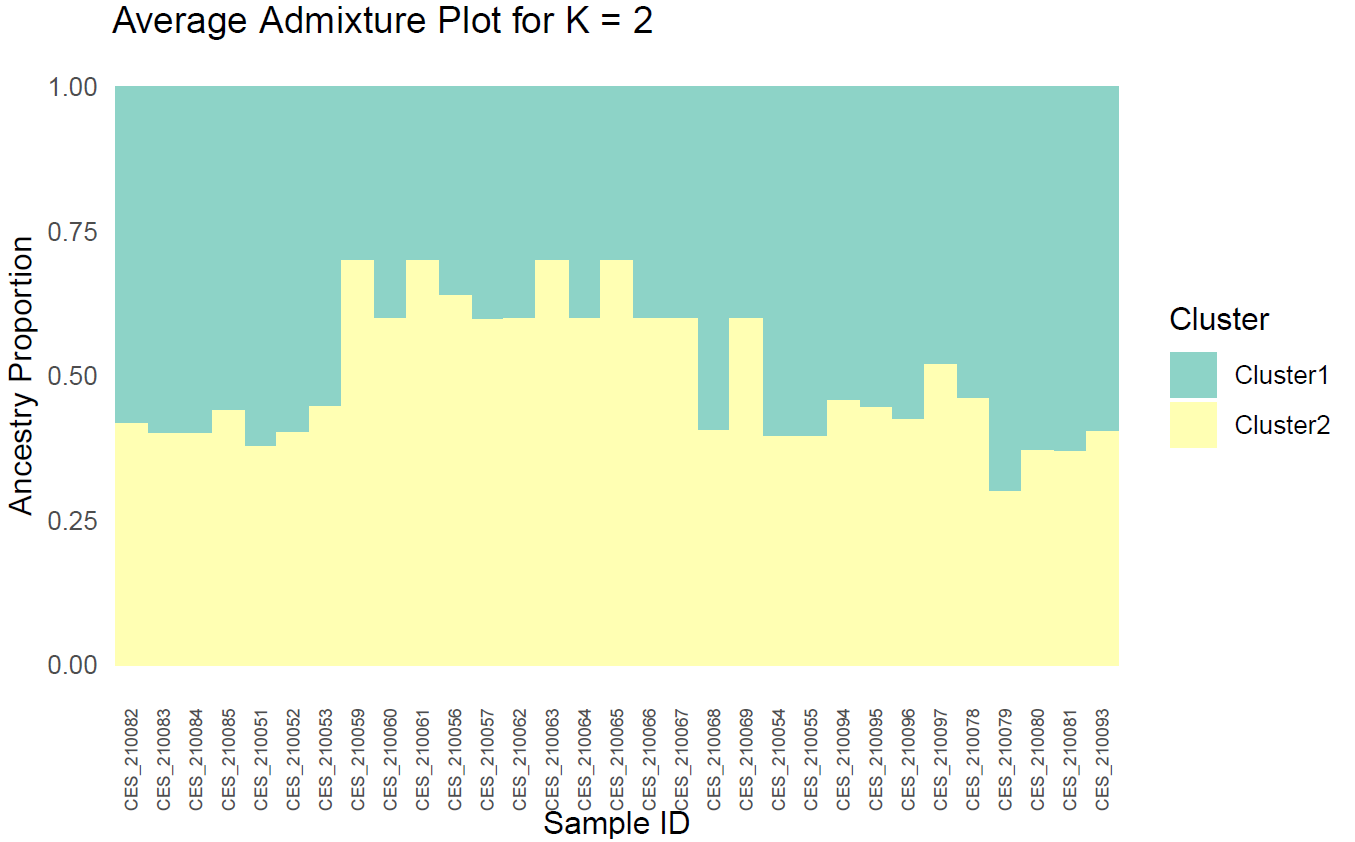

**Figure S3:** Admixture plot for K=4. Samples are plotted from north to south (left to right).

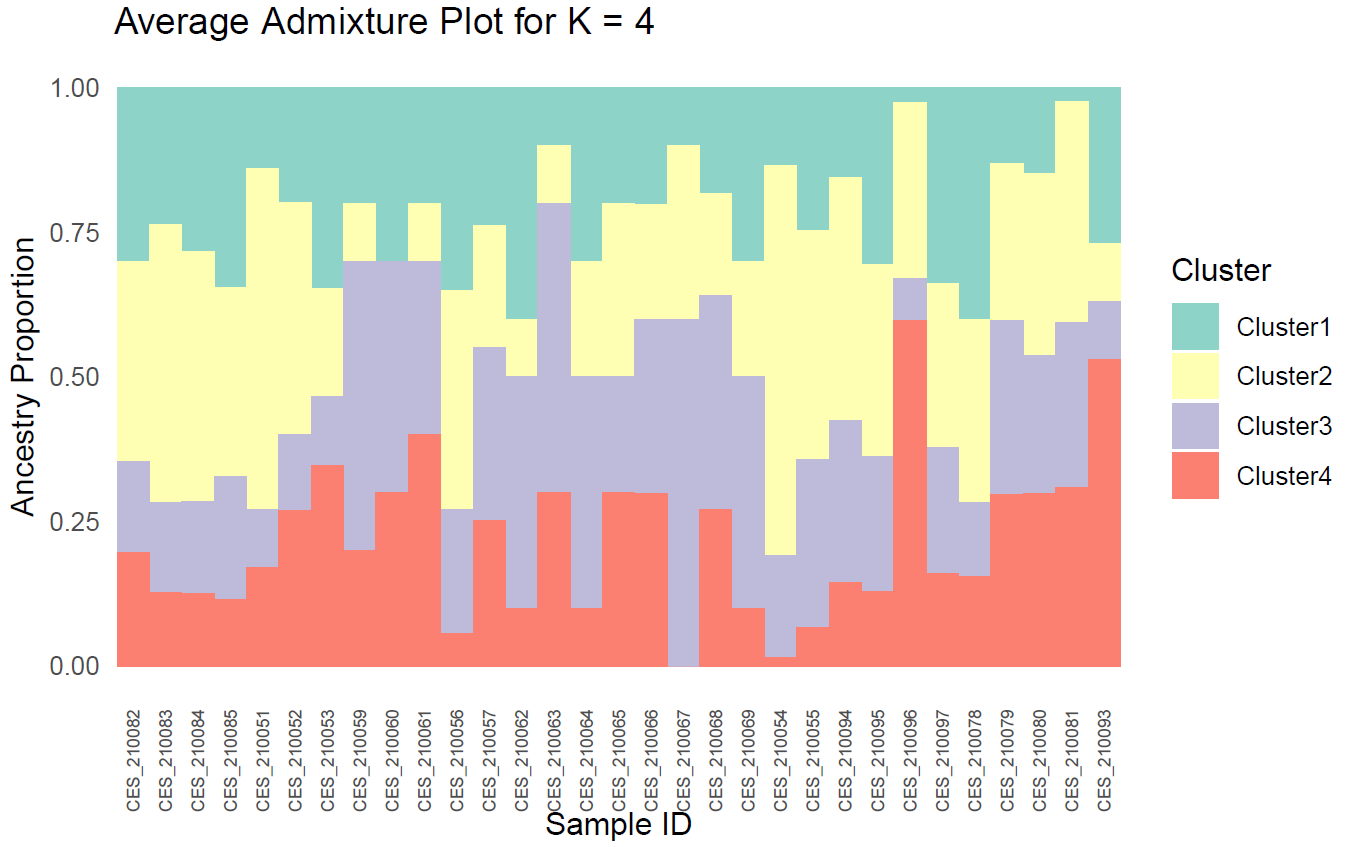

**Figure S4:** Heatmap of pairwise *F_ST_* comparisons between 11 populations

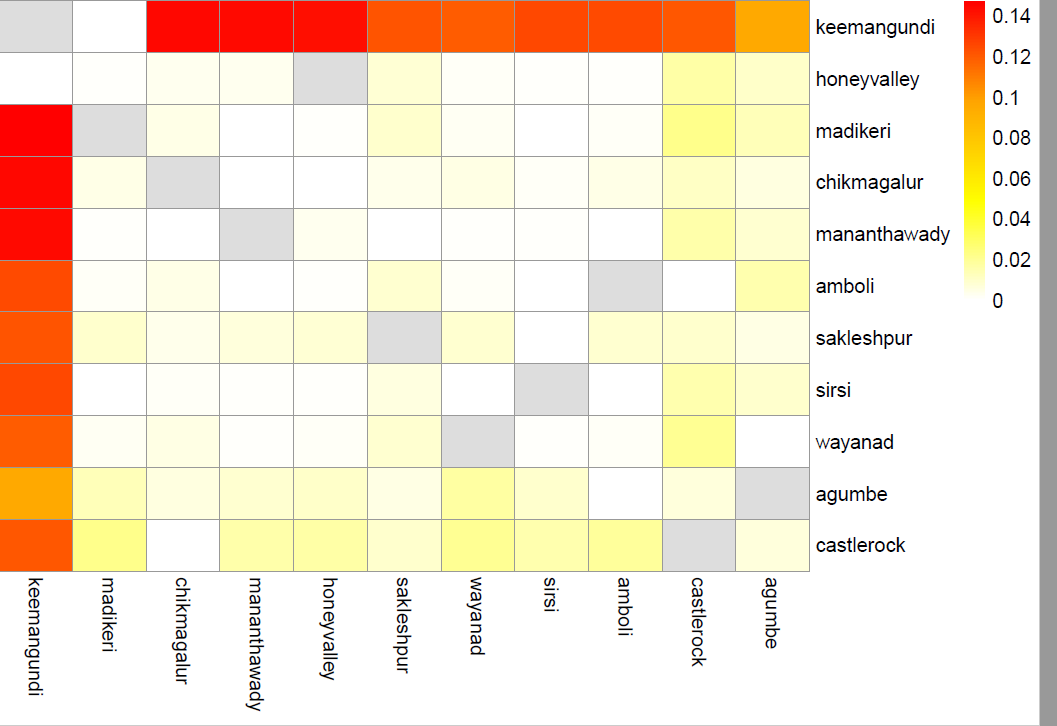

**Figure S5:** Line plots of He and F for the main dataset. Samples are plotted from south to north (left to right)

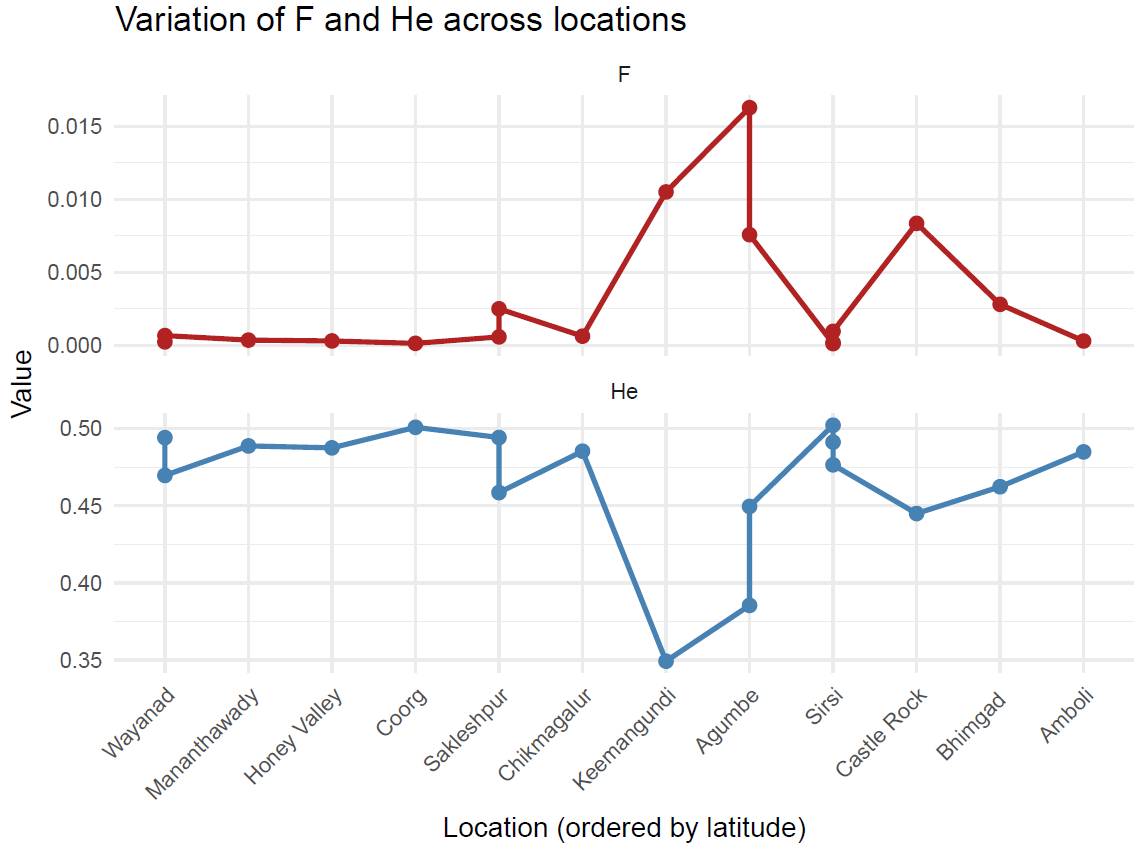

**Figure S6:** DHARMA diagnostics of the best model and residual structures

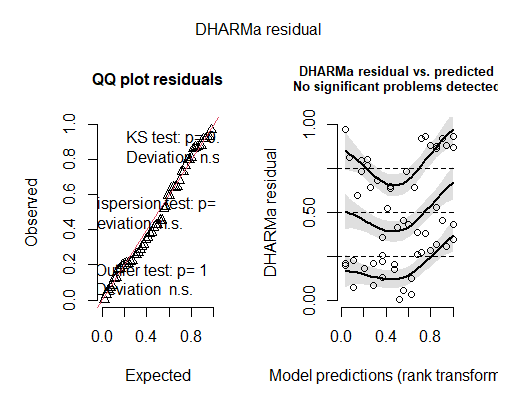

**Figure S7:** Historical demographic trajectory for the whole species

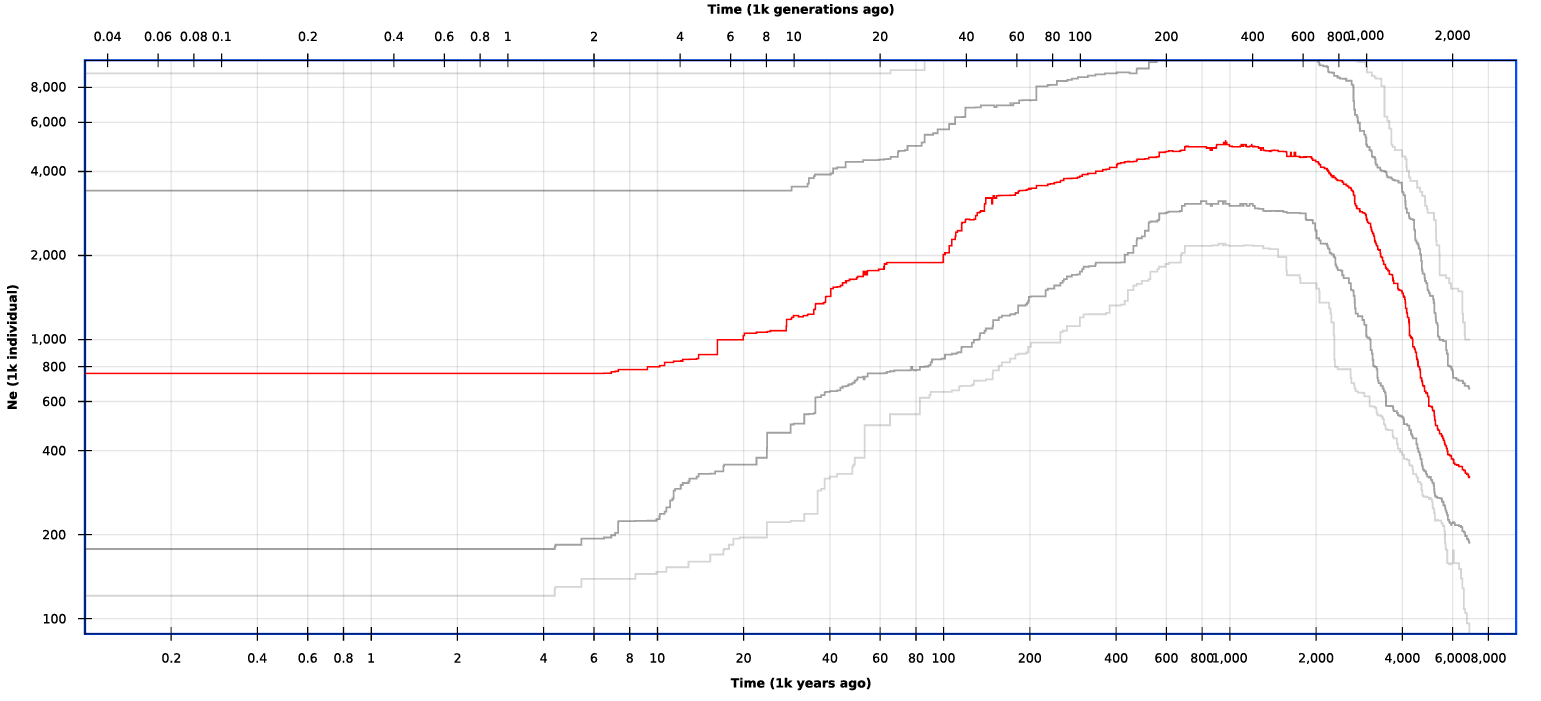

**Figure S8:** PCAngsd plot for the downsampled dataset. Samples are marked according to their location. Crosses indicate samples from the northern region, plus indicate samples from the southern part, and boxes denote samples from the central region.

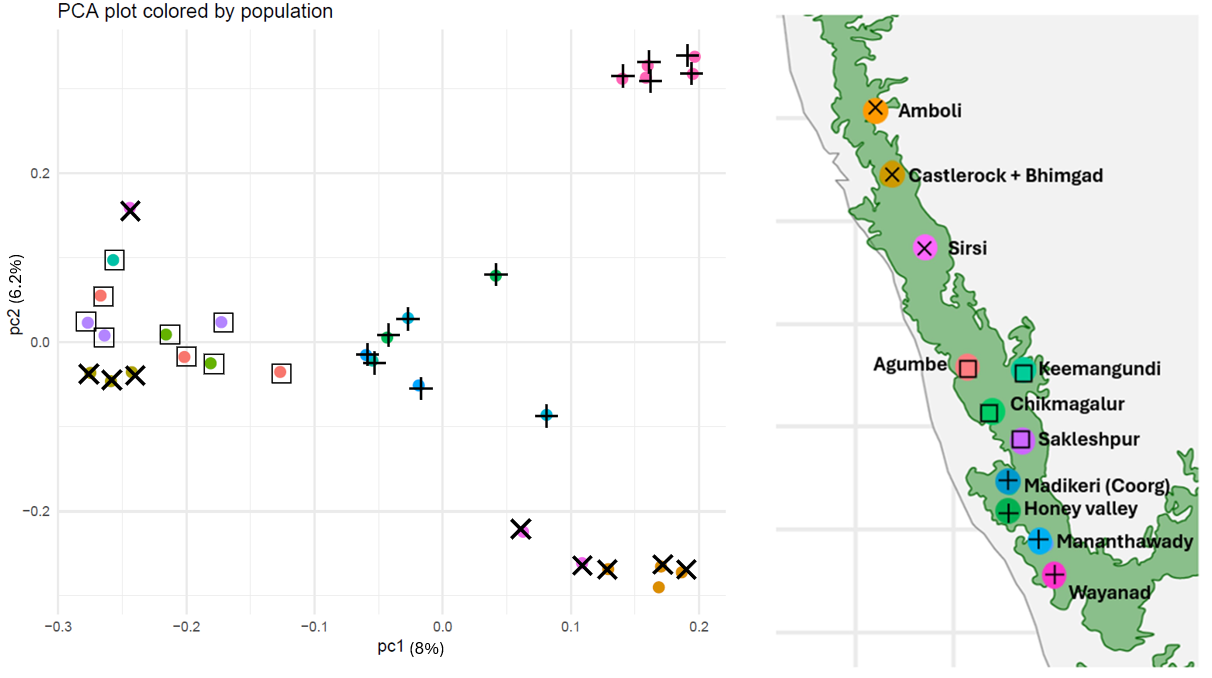

**Figure S9:** NGSadmix plot for the downsampled dataset along with ΔK and CV error plots for the best k. Samples are plotted from north to south (left to right)

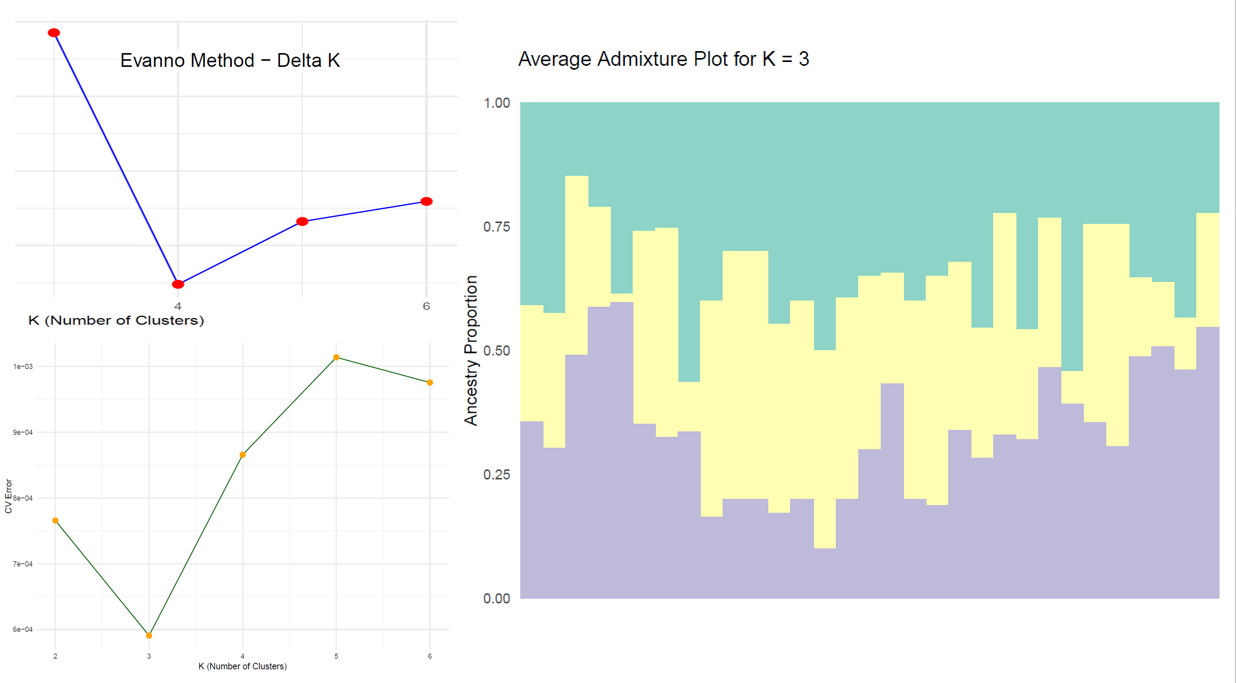

**Figure S10:** Proportion of heterozygous sites and inbreeding coefficient estimates from the downsampled dataset. Samples are plotted from south to north (left to right).

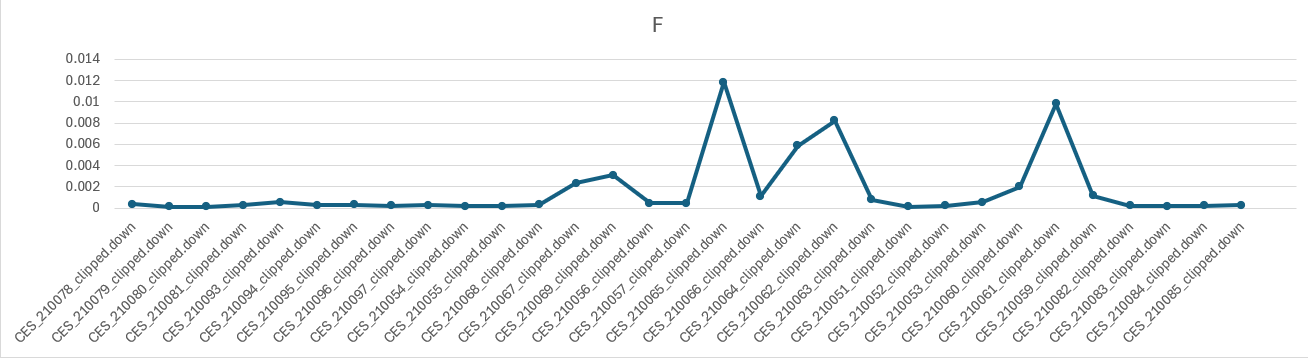

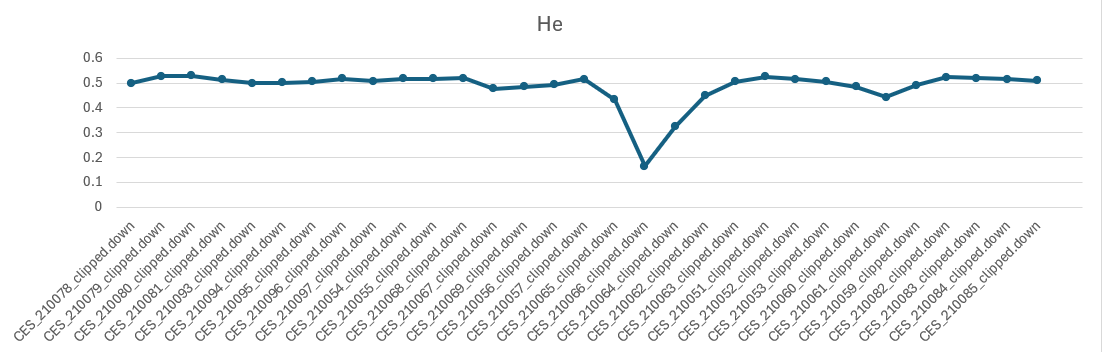
