## Supplementary figures and images for "Spatiotemporal variation in habitat suitability predicts genomic diversity and structure in a Western Ghats endemic Tarantula (*Thrigmopoeus truculentus*)"

### Supplementary material 3

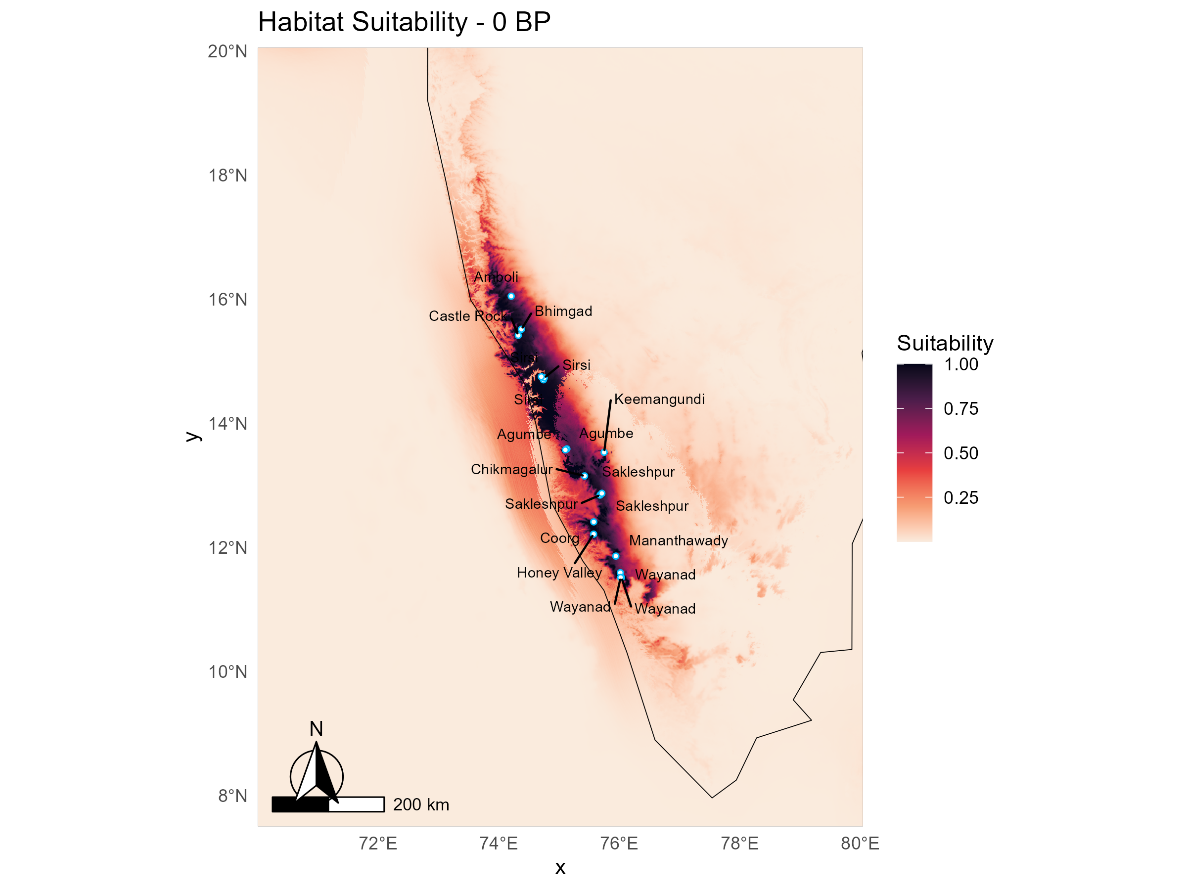

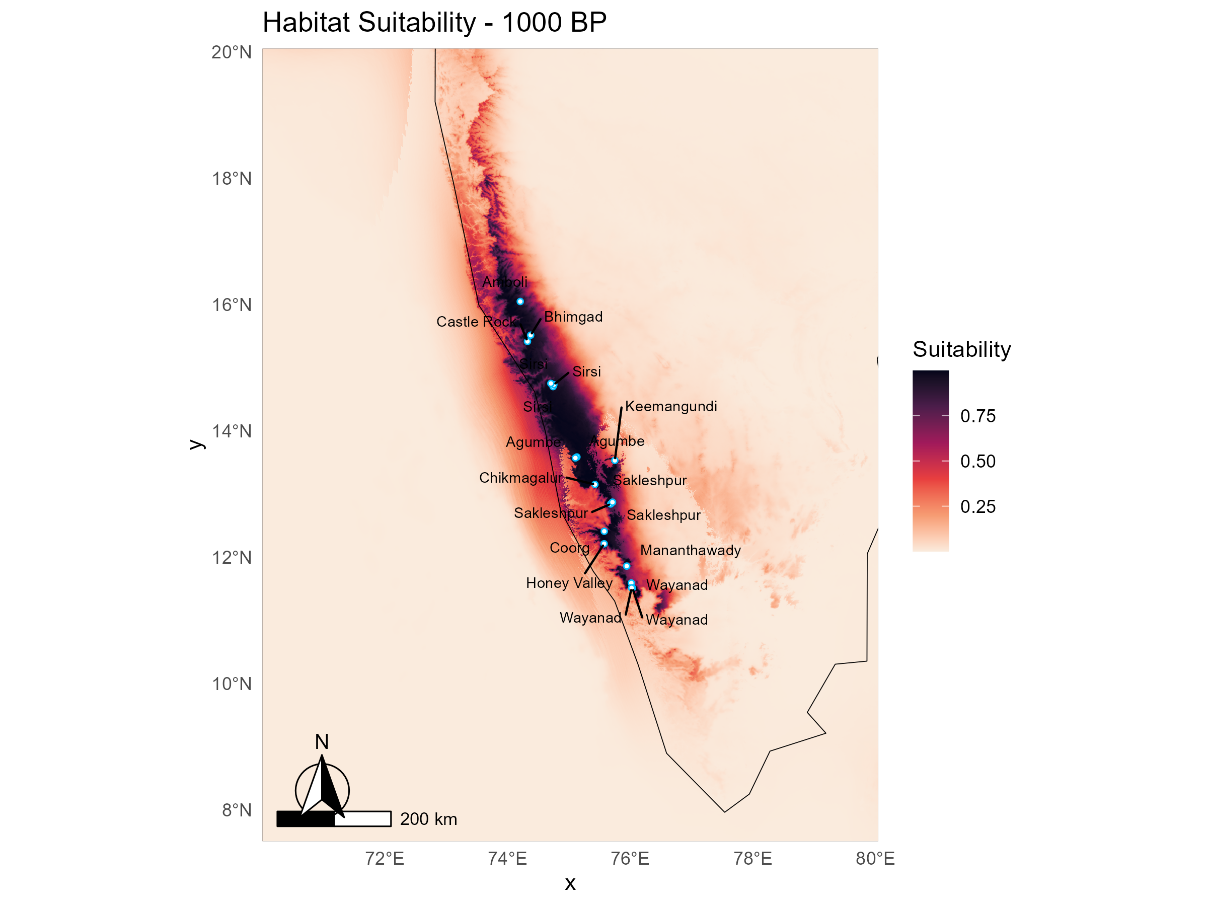

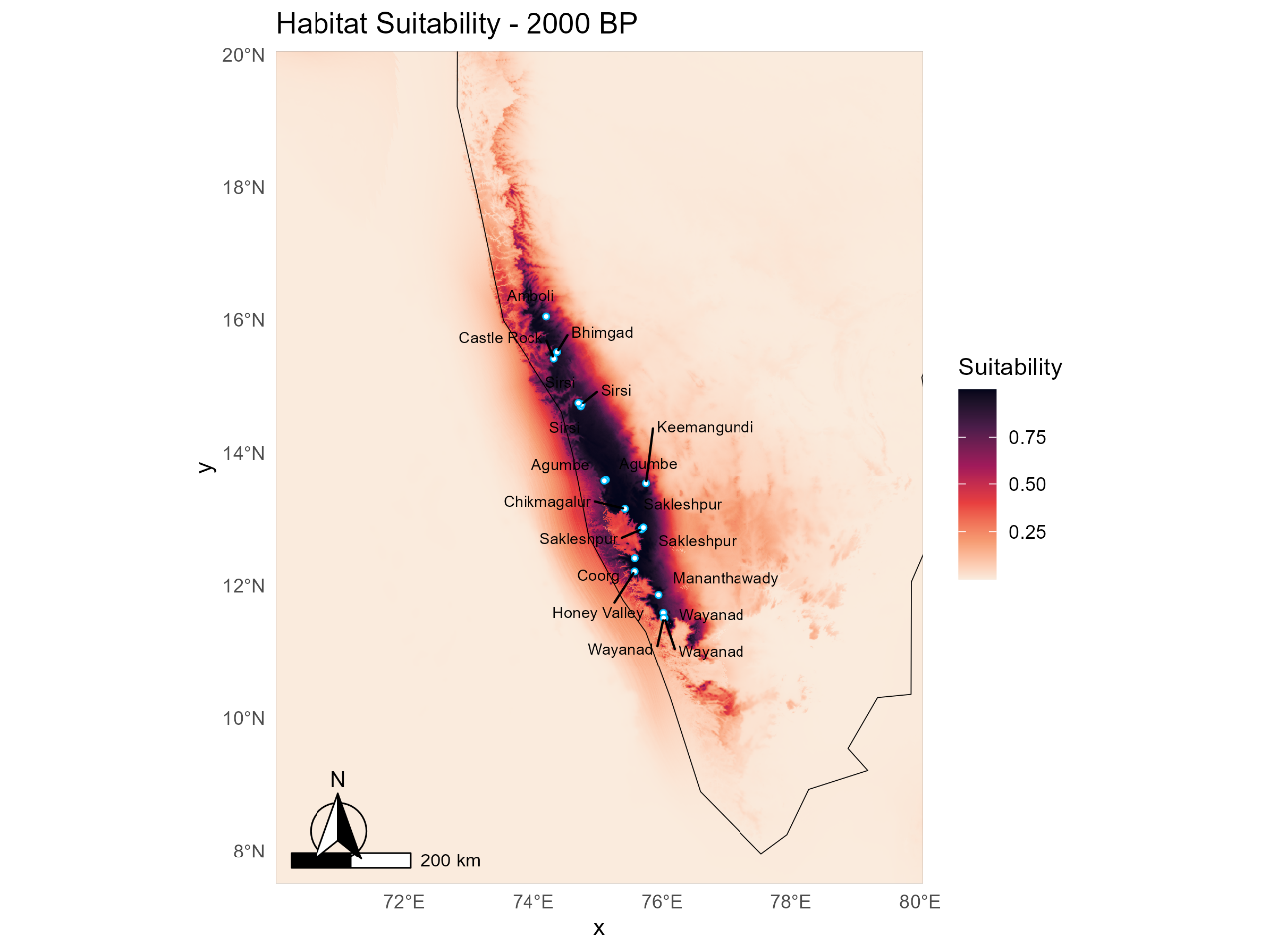

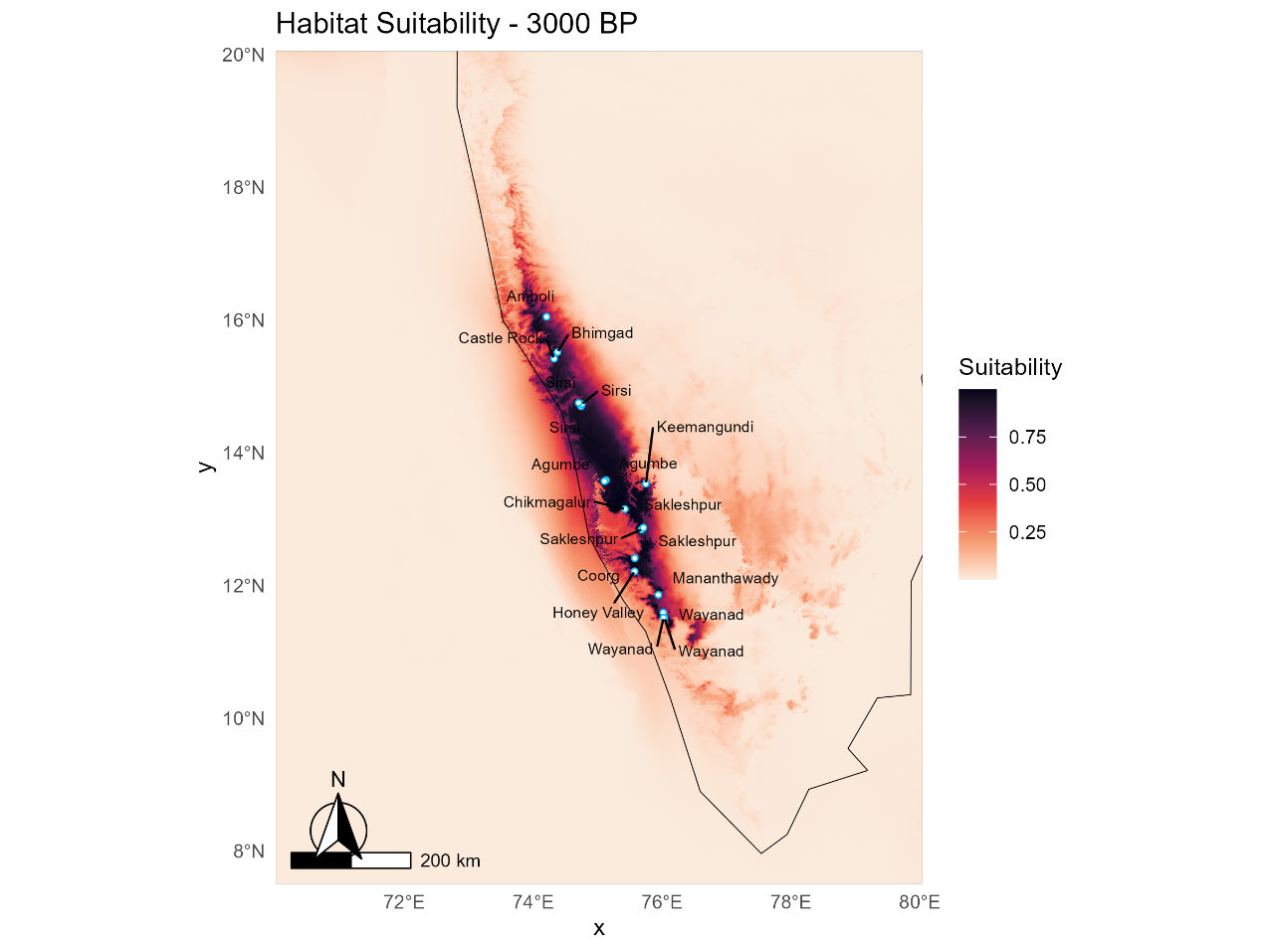

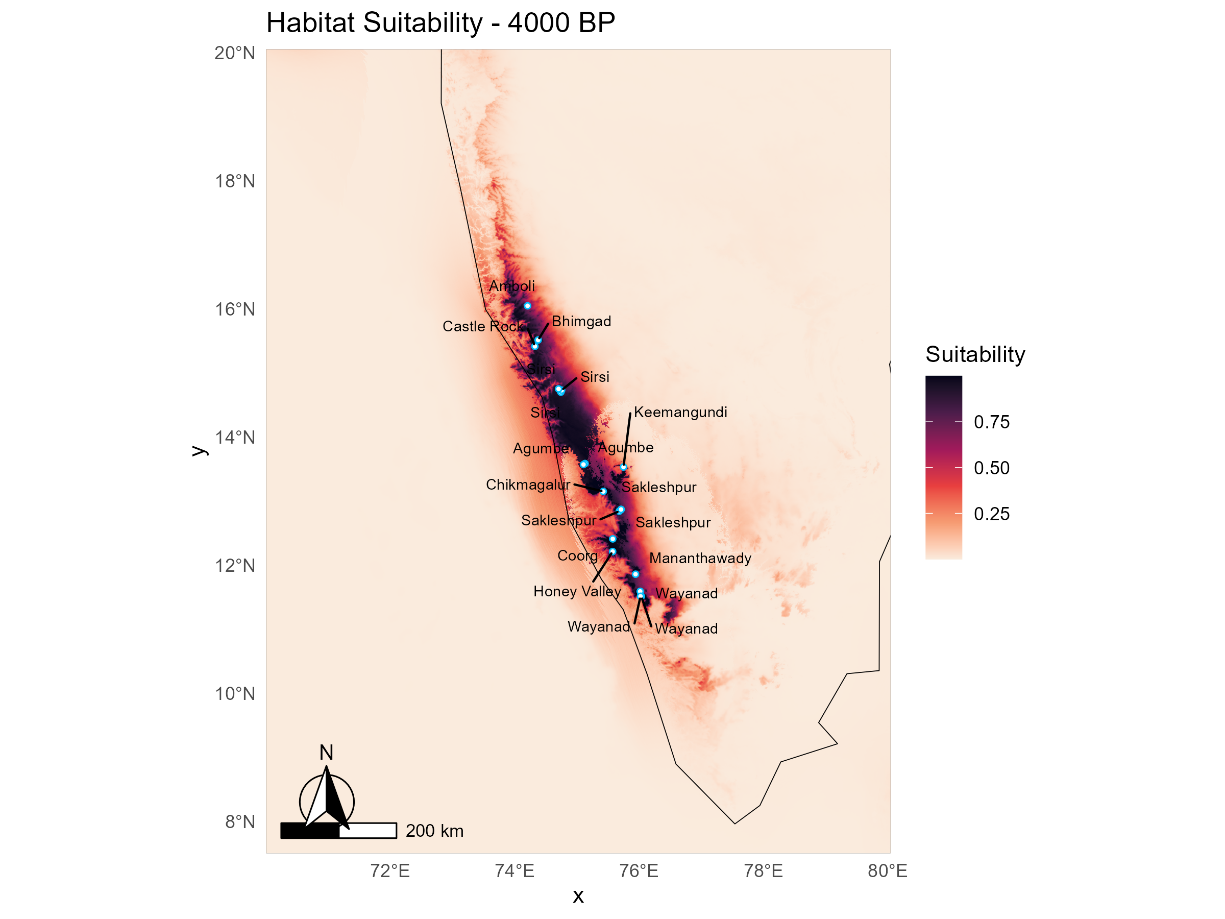

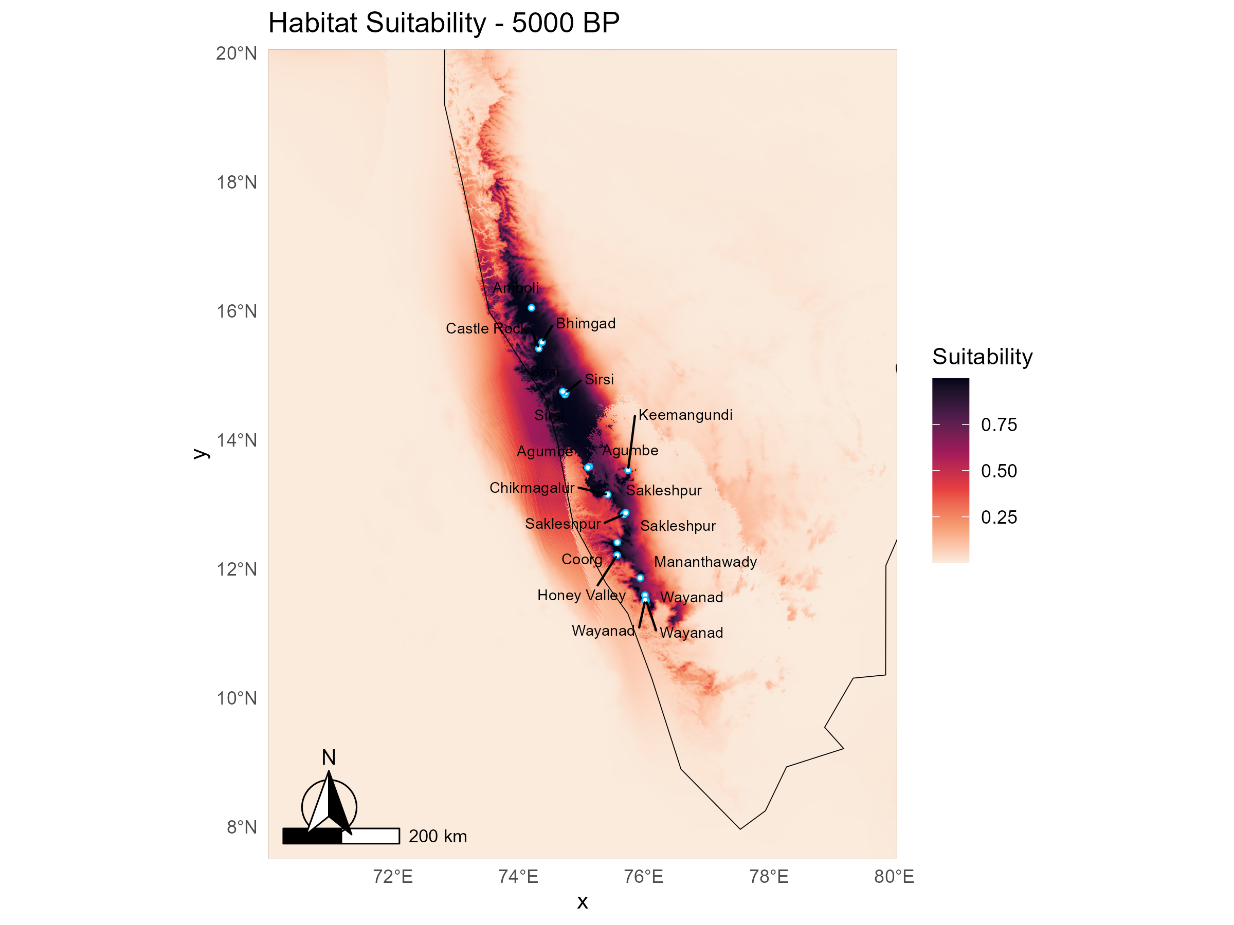

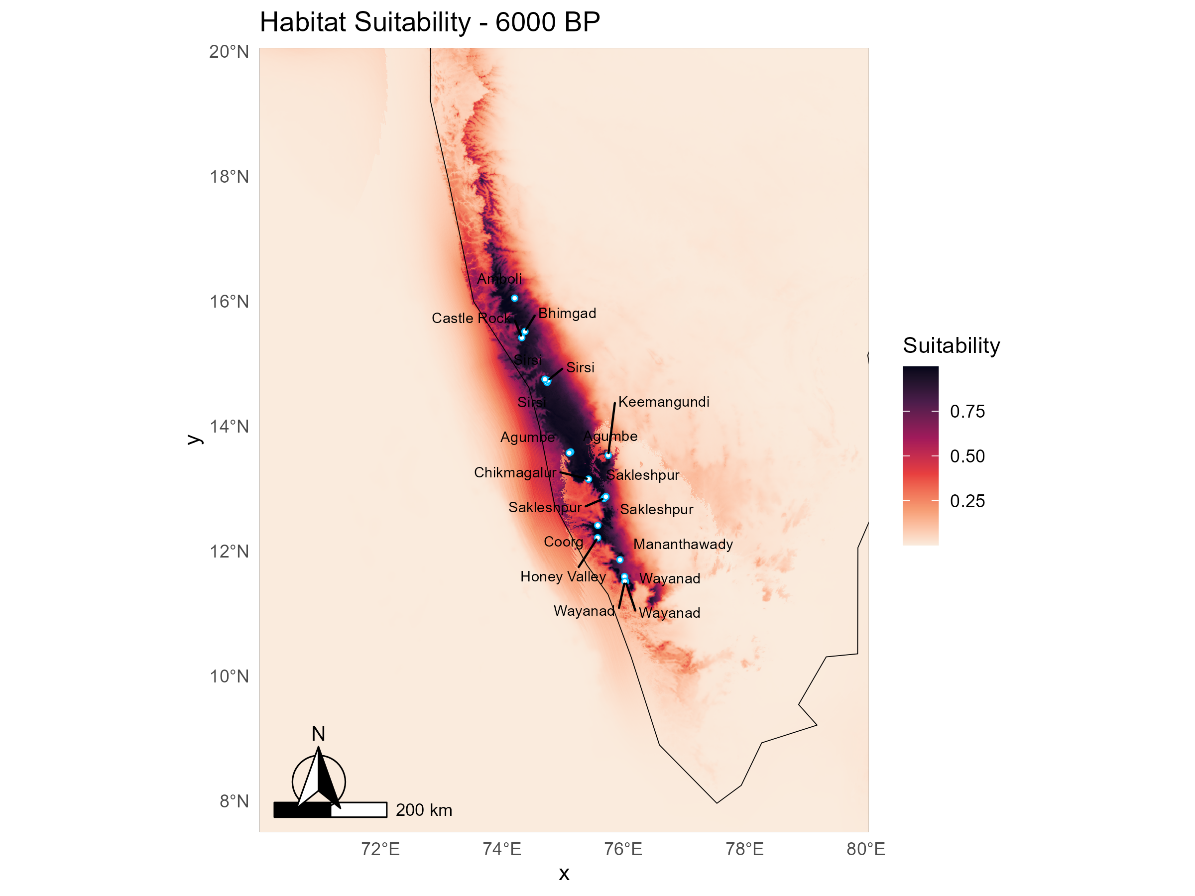

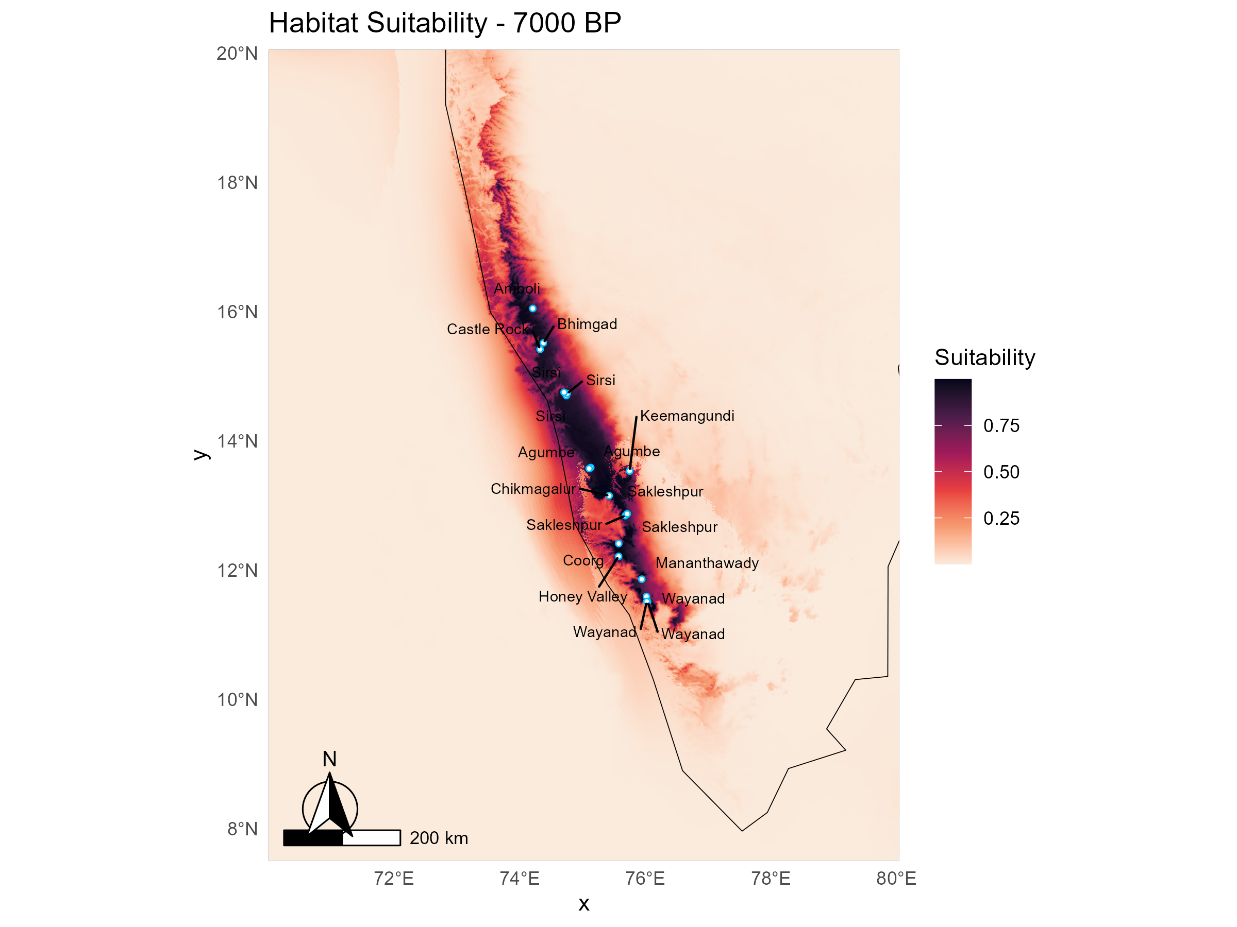

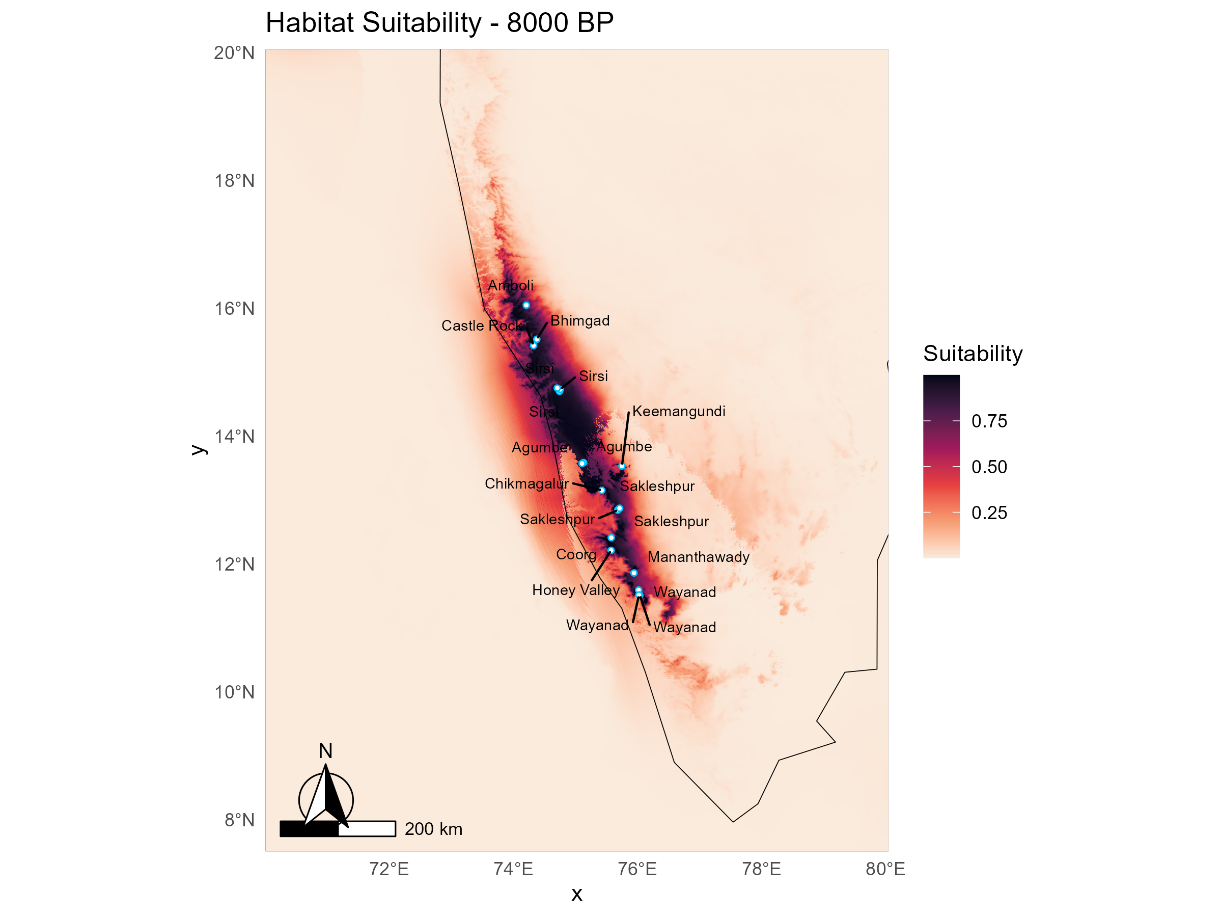

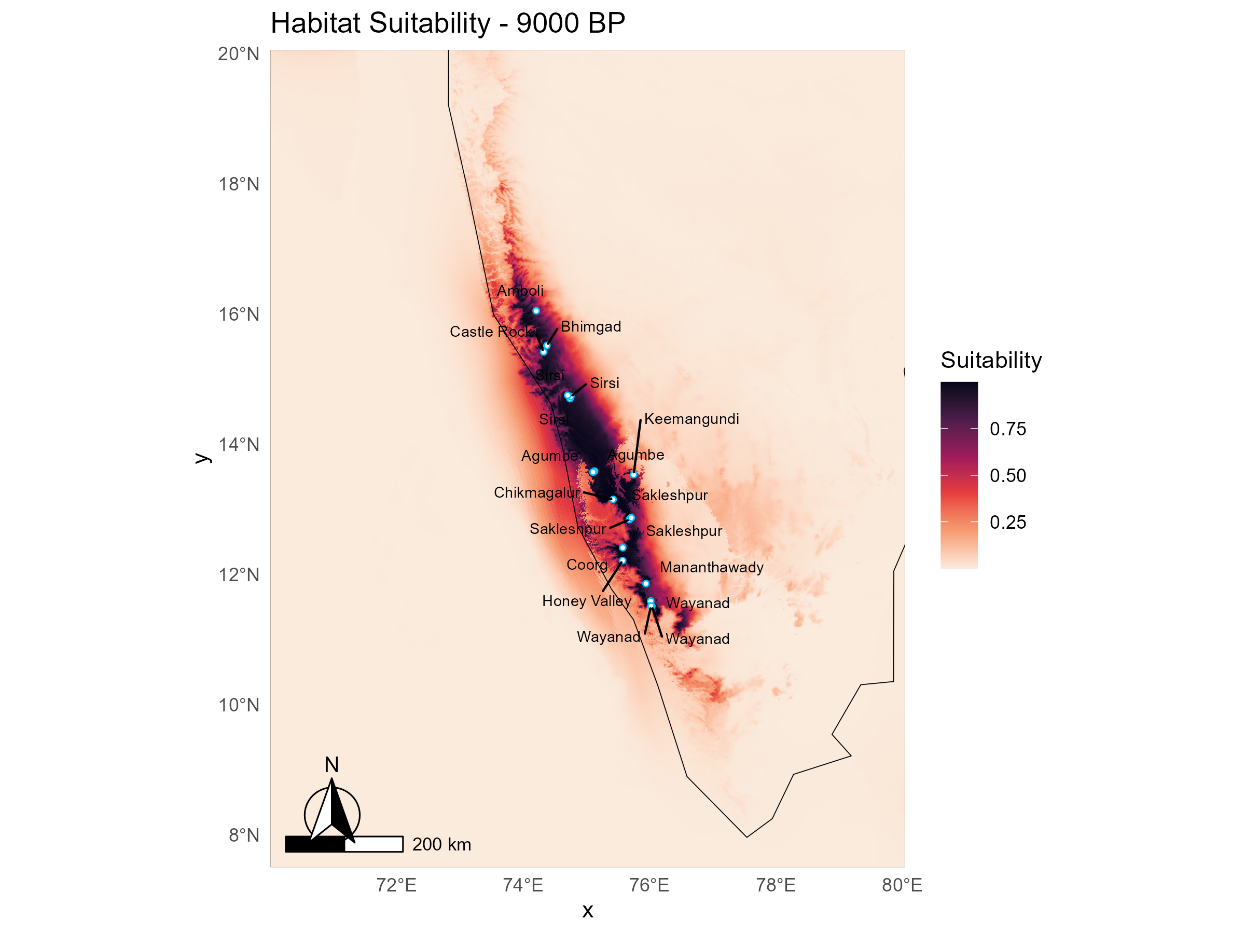

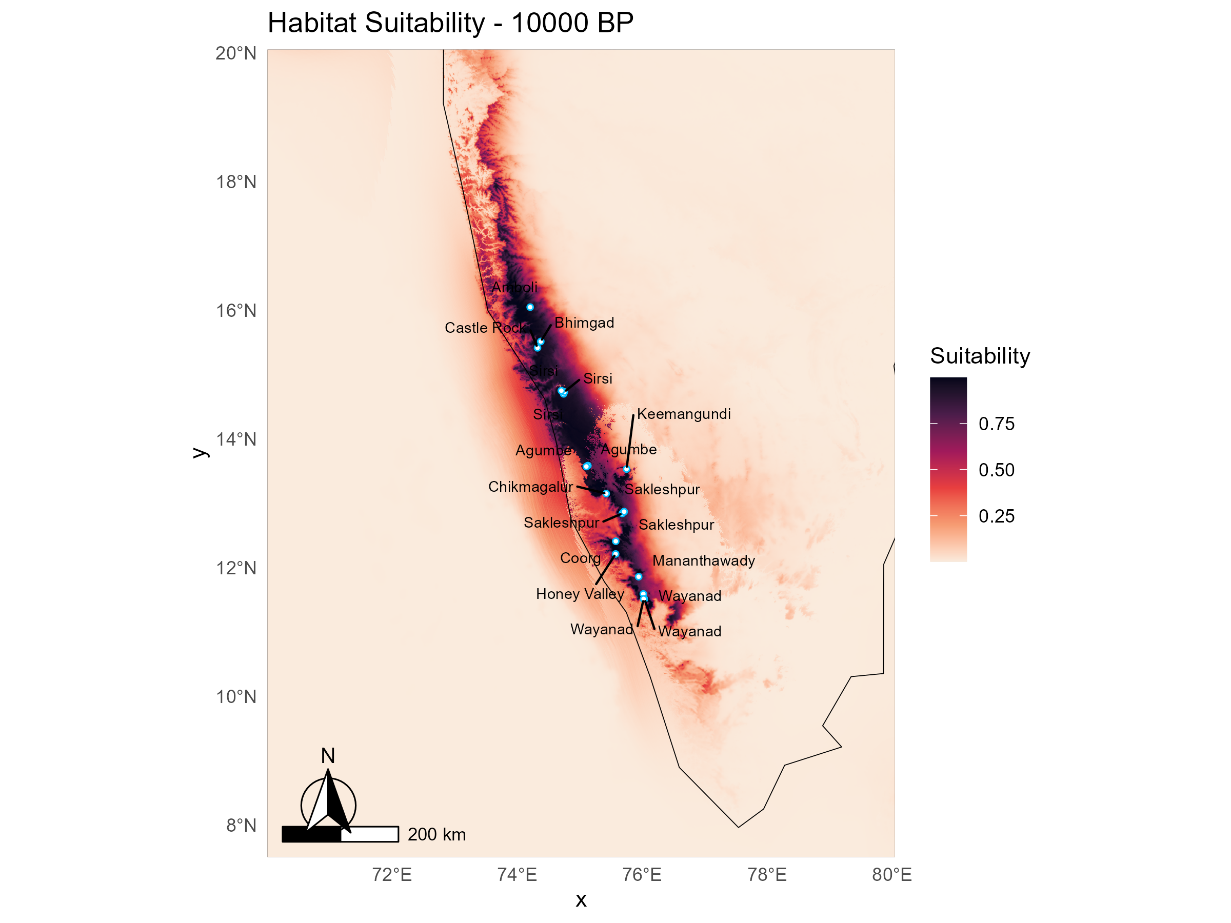

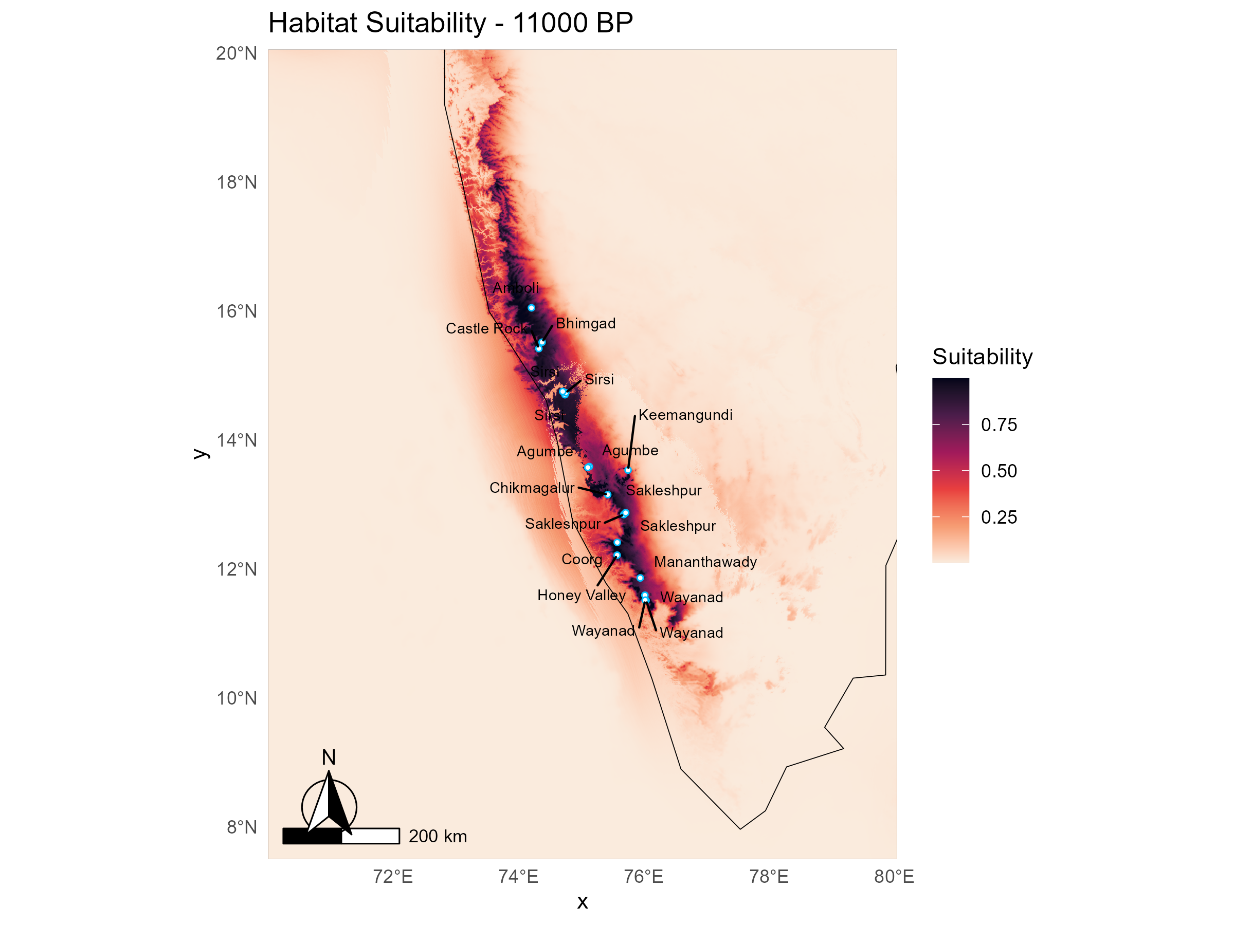

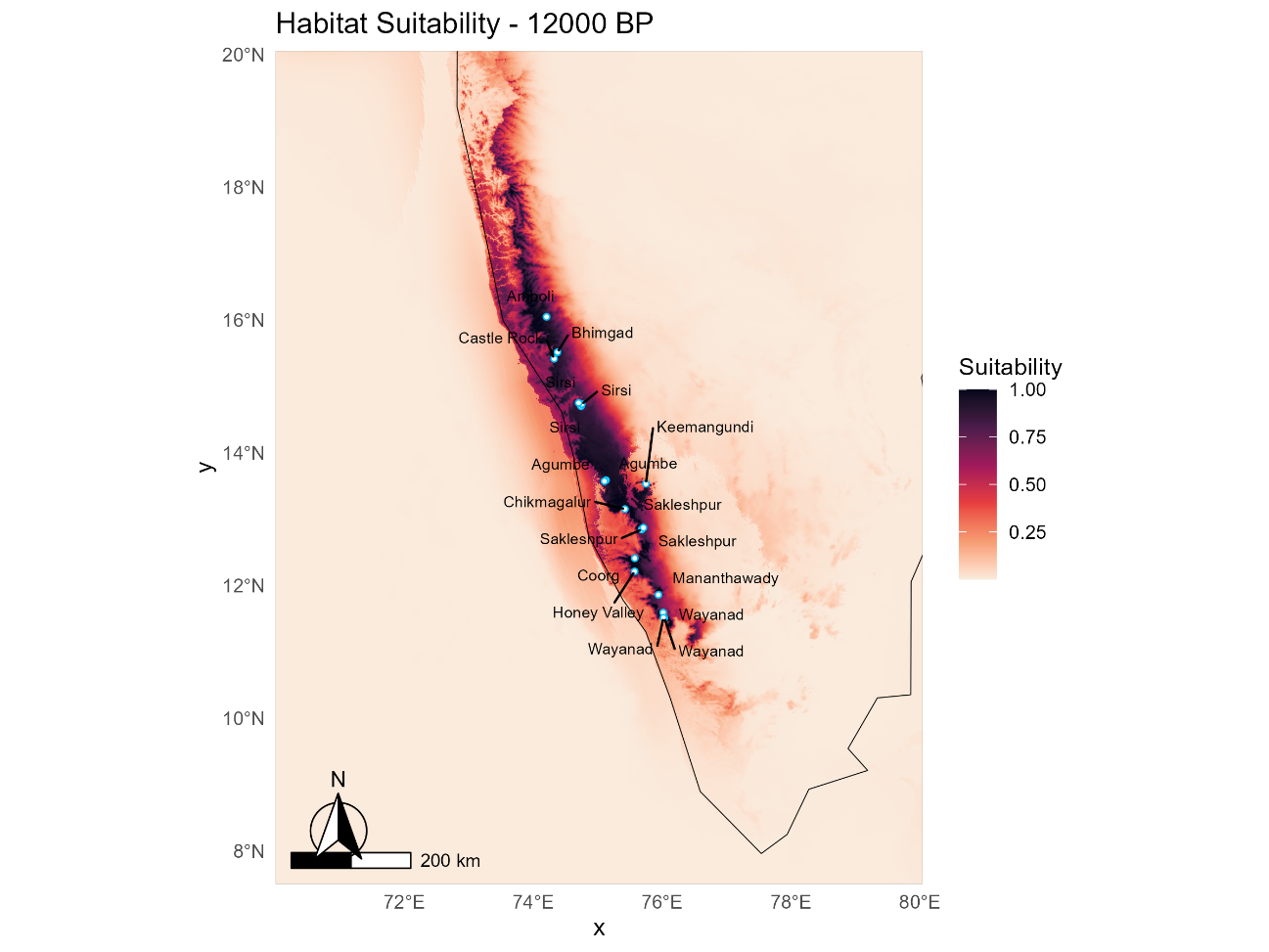

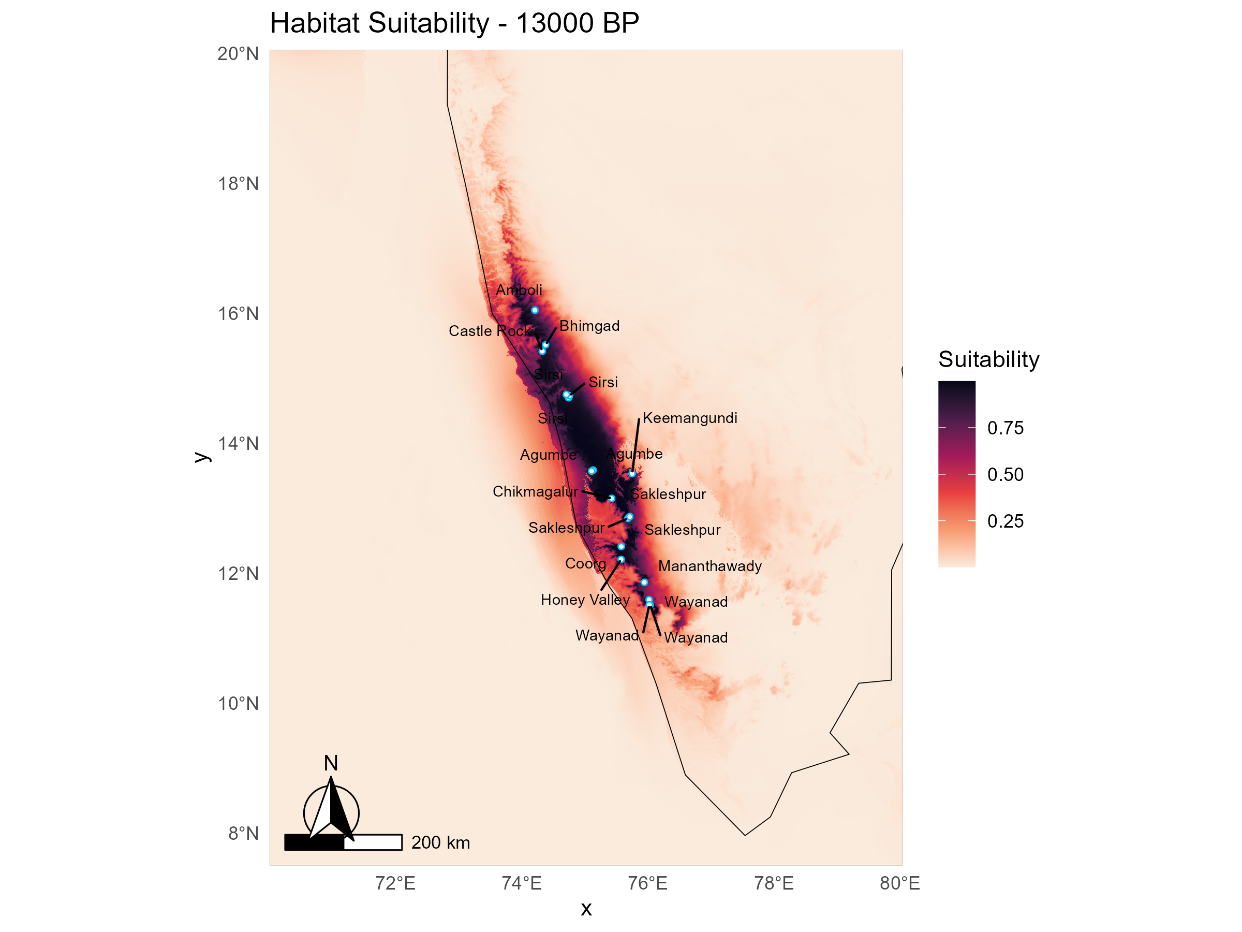

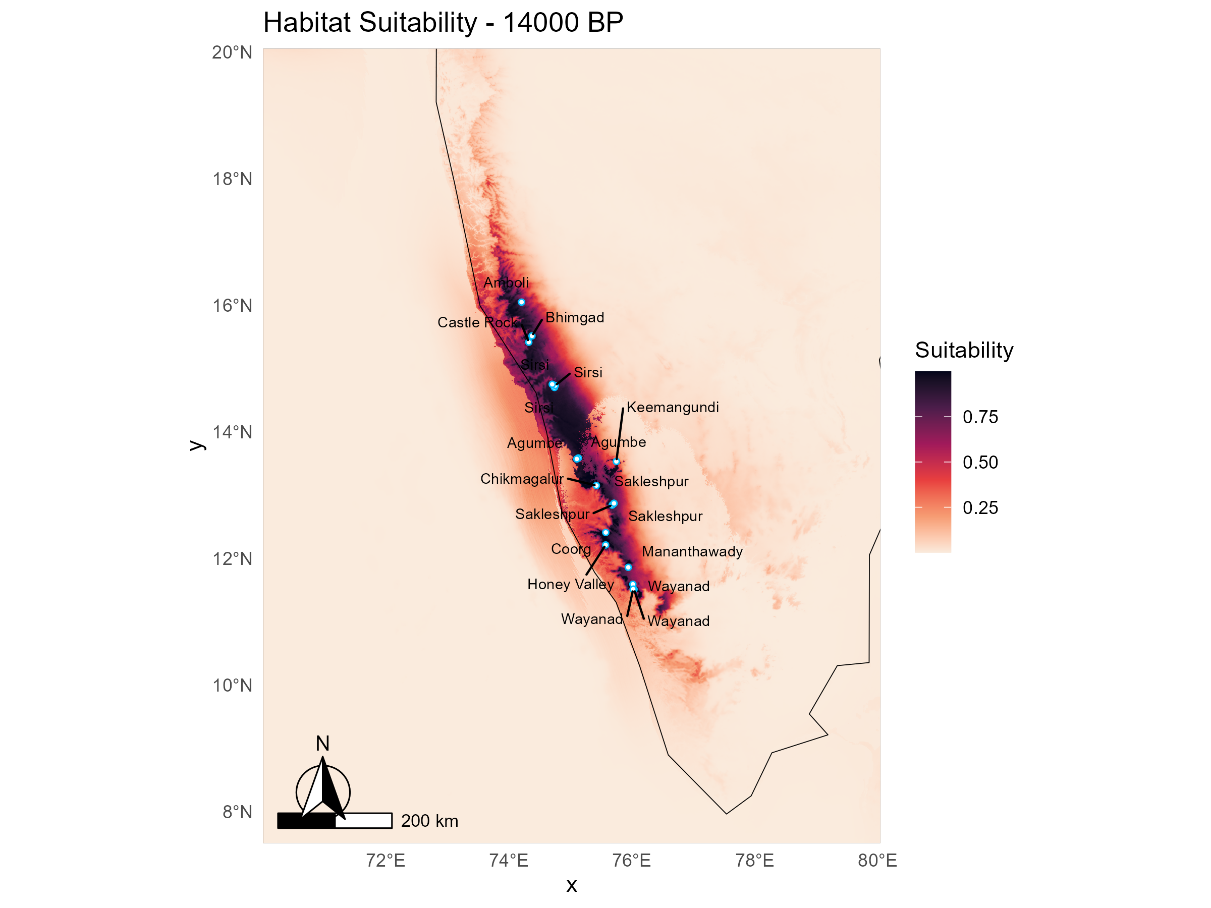

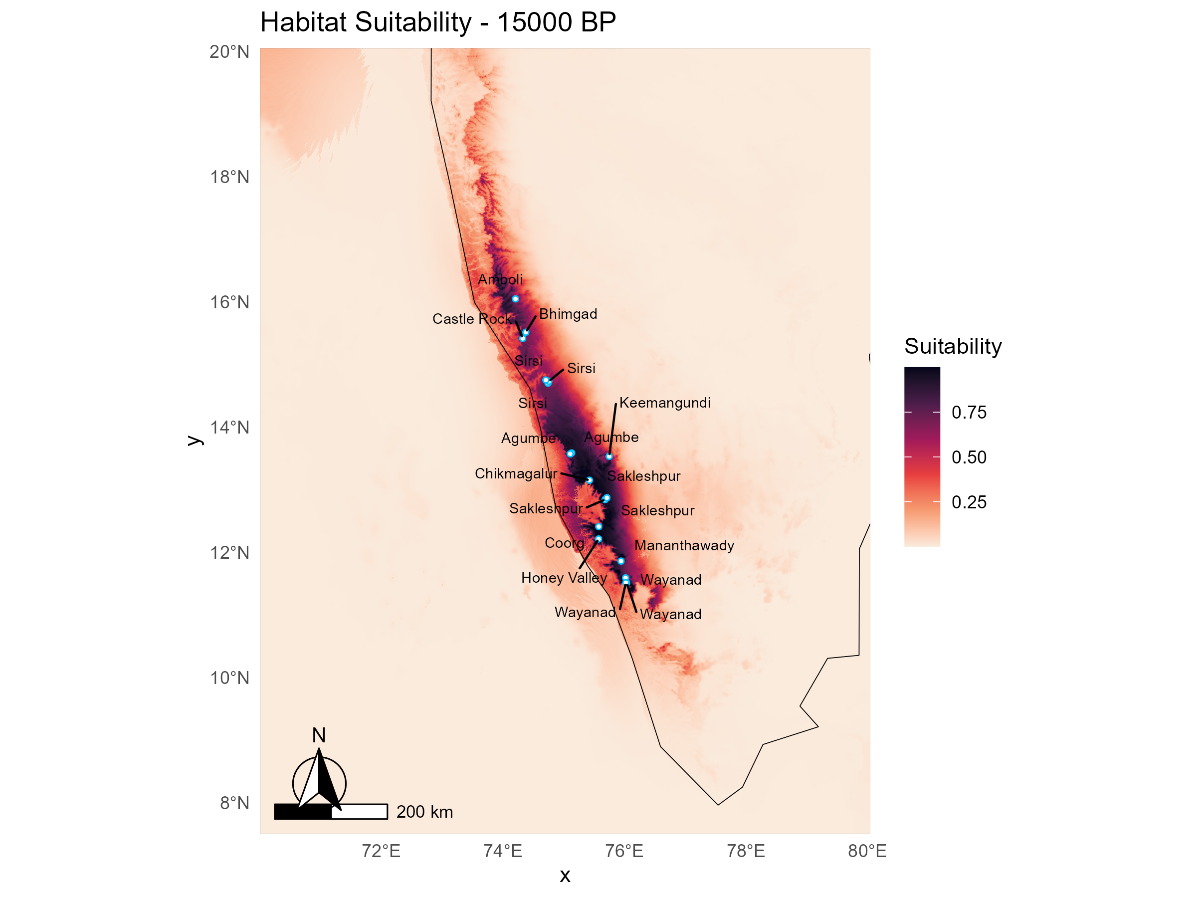

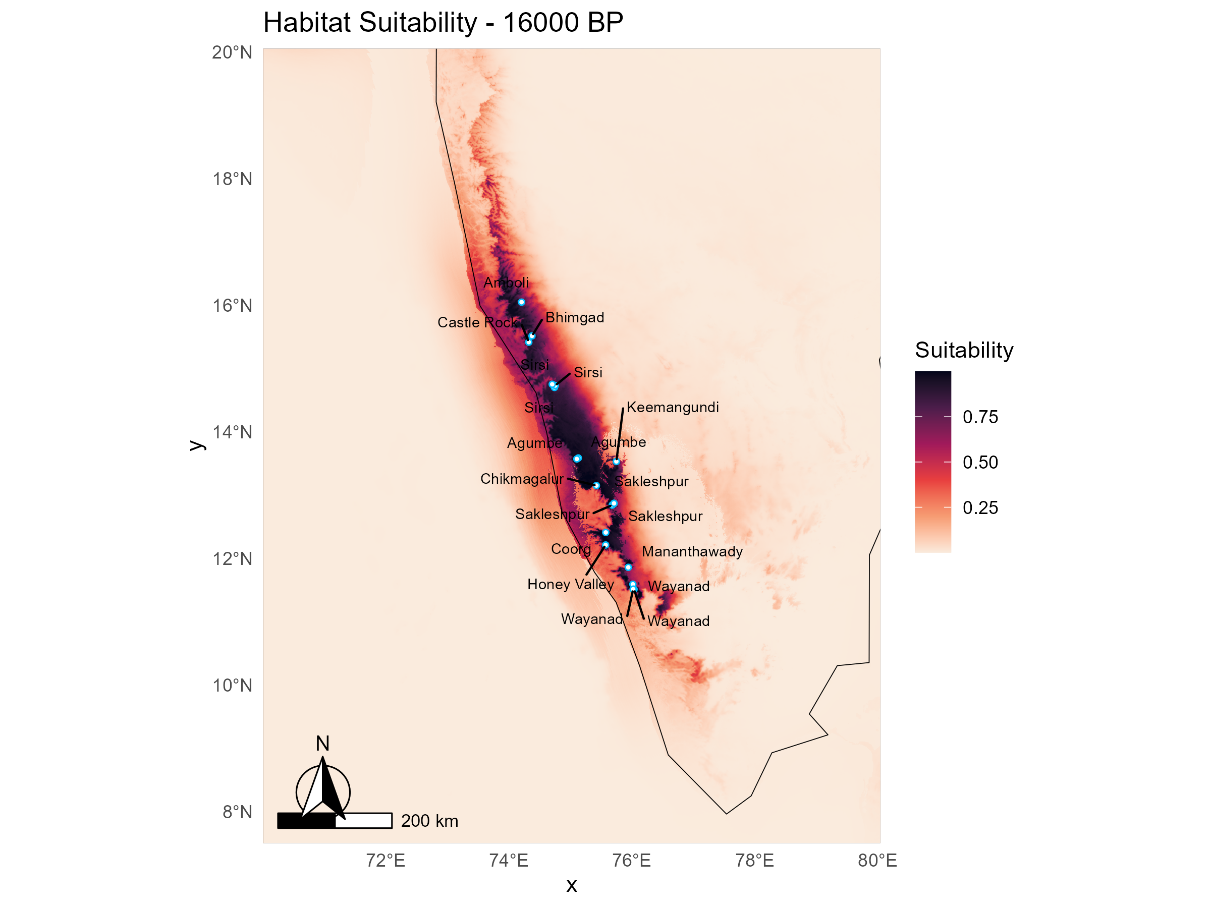

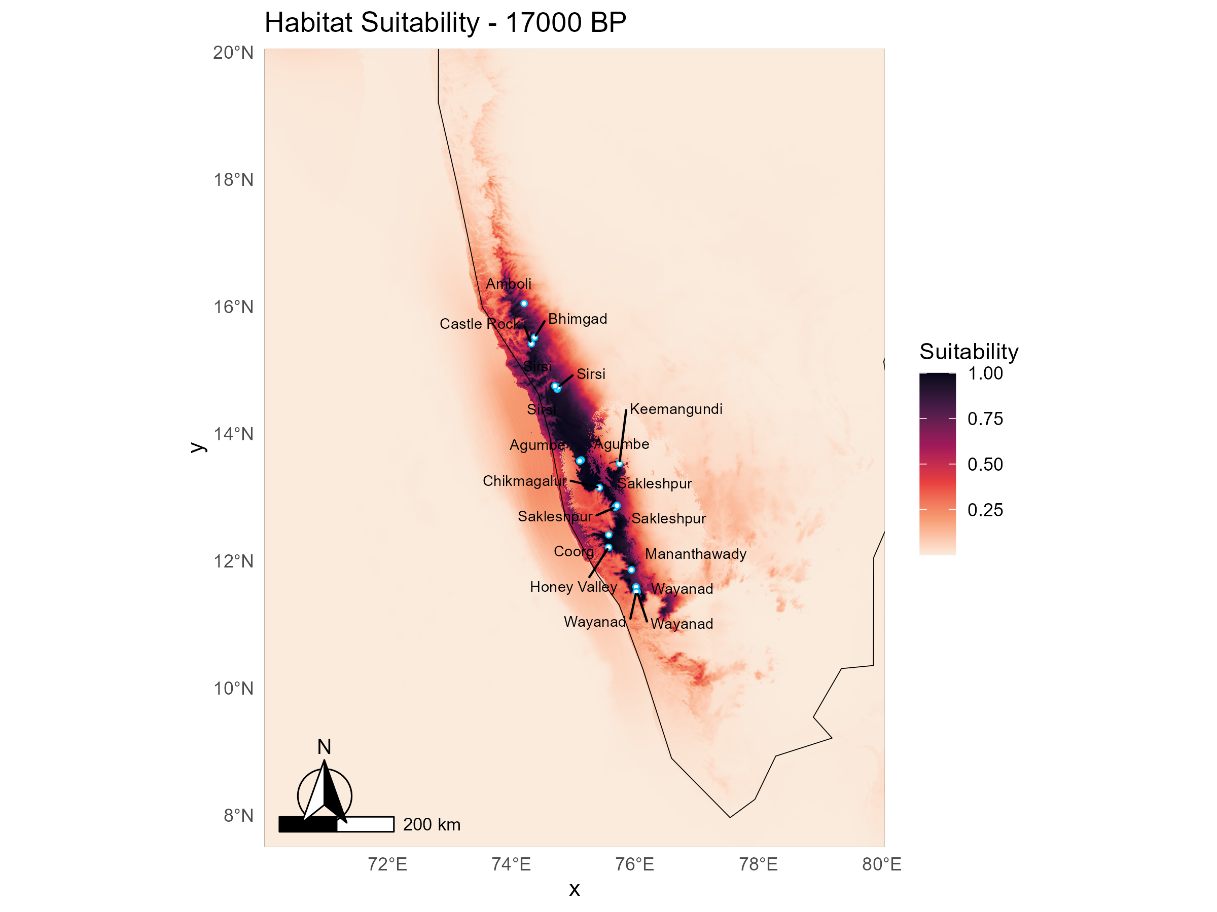

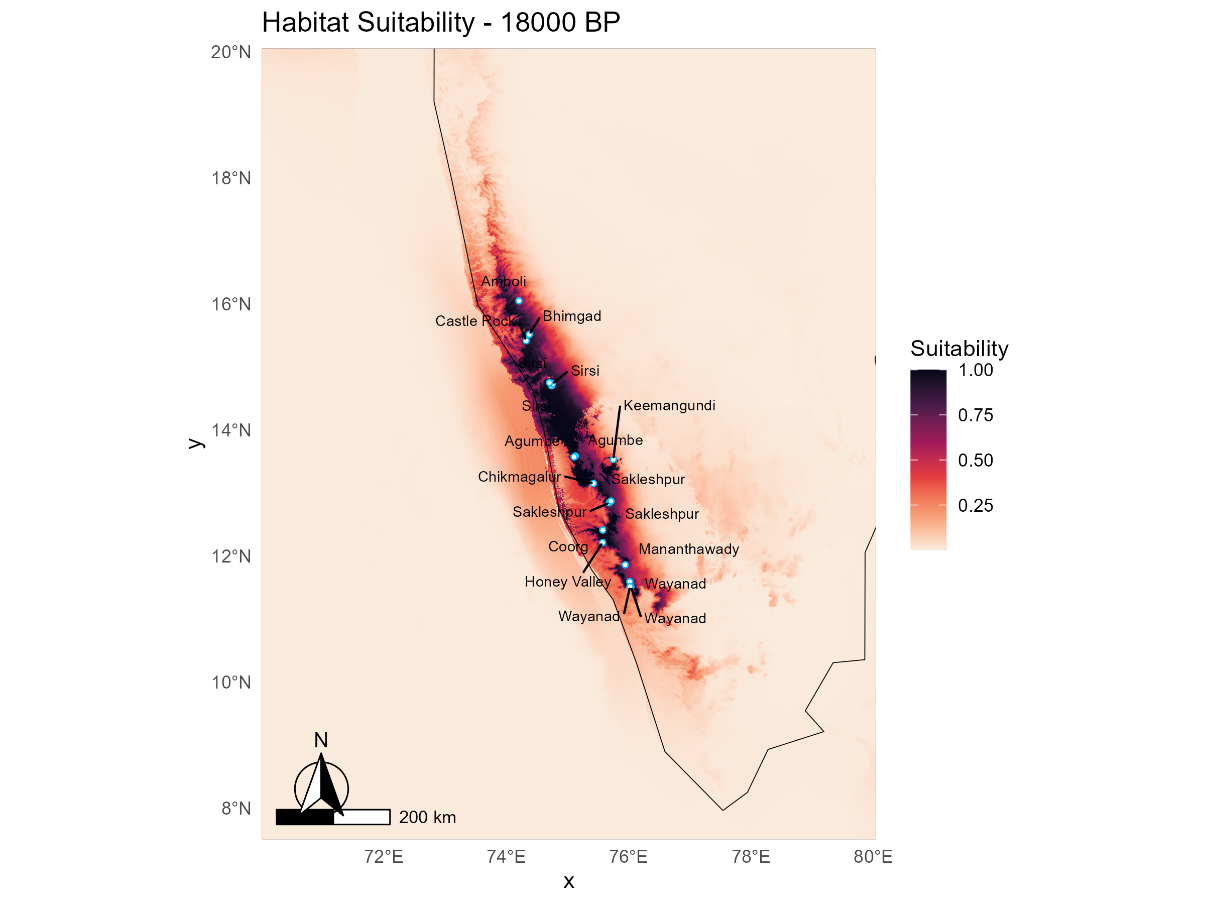

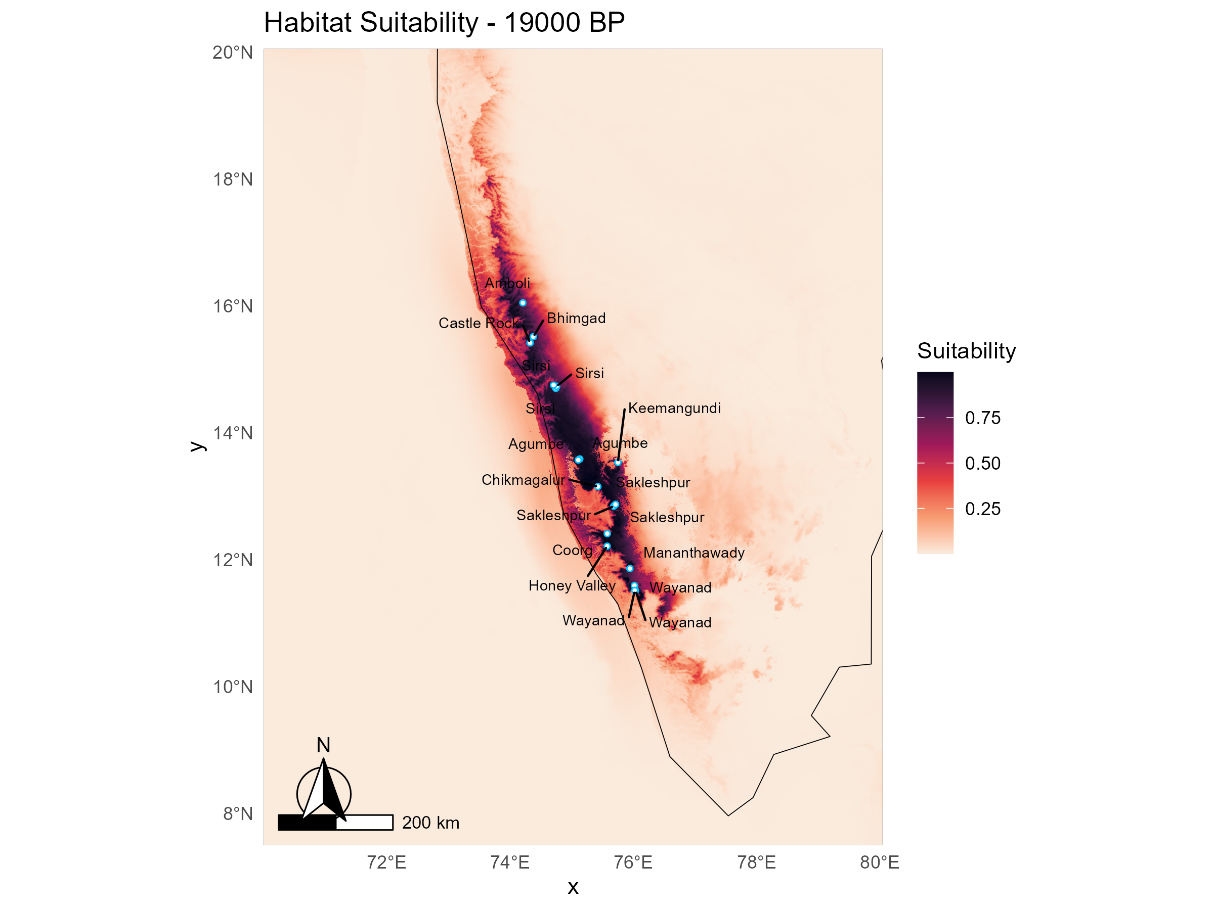

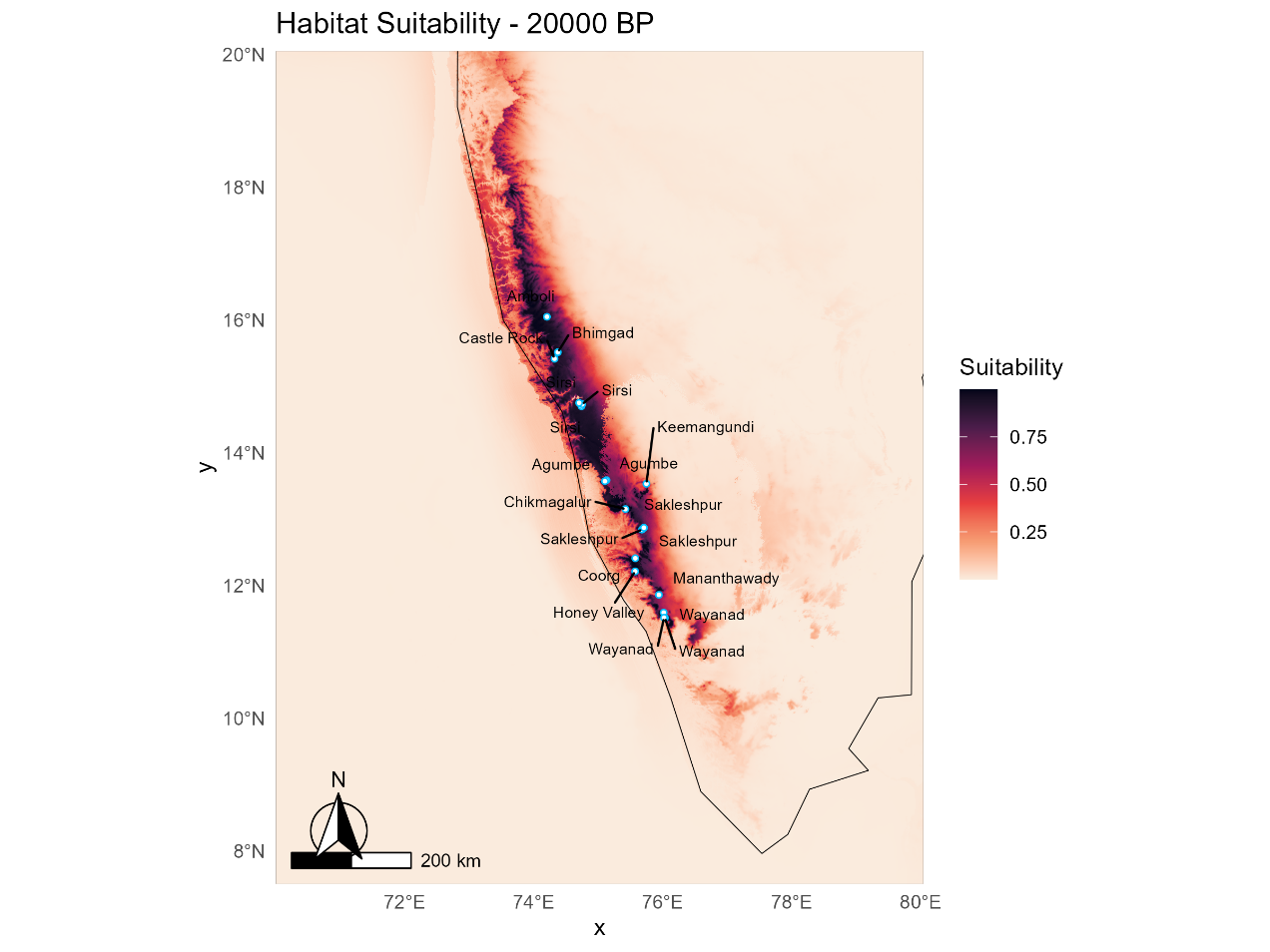
