## Supplementary material 1 for "Spatiotemporal variation in habitat suitability predicts genomic diversity and structure in a Western Ghats endemic Tarantula (*Thrigmopoeus truculentus*)"

**Section S1: Details of molecular and sequencing protocols**

**1. Genomic DNA Extraction, Qualitative and Quantitative analysis**

Genomic DNA was extracted from the received spider leg tissue samples using commercially available QIAamp DNA Mini Kit (Qiagen) as per manufacturer’s instruction. The quality and quantity of the extracted DNA sample was checked on NanoDrop followed by agarose gel electrophoresis.

**2. Preparation of 2 × 150 Shotgun library**

The QC passed and confirmed DNA samples were processed for paired-end sequencing library using QIAseq FX DNA Library Preparation Kit as per manufacturer’s instruction. Briefly, approximately 100ng of DNA was processed for fragmented, end-repaired and adaptor ligation in a single tube as per the kit protocol. The end-repaired DNA was processed for size selection using AMPure XP bead. The size-selected products were PCR amplified with the index primer as described in the kit protocol.

**3. Quantity and quality check (QC) of library on Agilent 4200 Tape Station**

The PCR enriched libraries were analyzed on 4200 Tape Station system (Agilent Technologies) using D1000 Screen tape as per manufacturer instructions.

**4. Cluster Generation and Sequencing**

After obtaining the Qubit concentration for the library and the mean peak size from Agilent Tape Station profile, the PE Illumina library was loaded onto Illumina for cluster generation and sequencing. Paired-End sequencing allows the template fragments to be sequenced in both the forward and reverse directions on Illumina. The kit reagents were used in binding of samples to complementary adapter oligoes on paired-end flow cell. The adapters were designed to allow selective cleavage of the forward strands after re-synthesis of the reverse strand during sequencing. The copied reverse strand was used to sequence from the opposite end of the fragment.

**Table S1: Raw read Statistics Per Sample**

| Sl no. | Sample ID | No. of PE reads | Data in GB | Average coverage |
| --- | --- | --- | --- | --- |
| 1 | CES 210051 | 13540543 | 4.09 | 0.1185x |
| 2 | CES 210052 | 11673397 | 3.53 | 0.1063x |
| 3 | CES 210053 | 12702026 | 2.12 | 0.068x |
| 4 | CES 210054 | 11341987 | 3.43 | 0.1118x |
| 5 | CES 210055 | 12719134 | 3.84 | 0.1134x |
| 6 | CES 210056 | 12161874 | 3.67 | 0.0697x |
| 7 | CES 210057 | 12072682 | 3.6 | 0.0591x |
| 8 | CES 210059 | 11499434 | 3.4 | 0.0344x |
| 9 | CES 210060 | 13685949 | 4 | 0.0322x |
| 10 | CES 210061 | 8802128 | 2.6 | 0.0309x |
| 11 | CES 210062 | 8028915 | 2.4 | 0.0213x |
| 12 | CES 210063 | 10720819 | 3.2 | 0.0353x |
| 13 | CES 210064 | 14742966 | 4.4 | 0.0358x |
| 14 | CES 210065 | 14020663 | 4.2 | 0.0418x |
| 15 | CES 210066 | 14028030 | 4.2 | 0.0455x |
| 16 | CES 210067 | 12916462 | 3.8 | 0.038x |
| 17 | CES 210068 | 9595071 | 2.8 | 0.0615x |
| 18 | CES 210069 | 23764980 | 7 | 0.0678x |
| 19 | CES 210078 | 10831484 | 3.27 | 0.0952x |
| 20 | CES 210079 | 15111040 | 4.56 | 0.1319x |
| 21 | CES 210080 | 12958567 | 3.91 | 0.1157x |
| 22 | CES 210081 | 11865265 | 3.58 | 0.1041x |
| 23 | CES 210082 | 12117761 | 3.66 | 0.1036x |
| 24 | CES 210083 | 12394977 | 3.74 | 0.1135x |
| 25 | CES 210084 | 12236474 | 3.70 | 0.1135x |
| 26 | CES 210085 | 8969036 | 2.71 | 0.0854x |
| 27 | CES 210093 | 10360431 | 3.13 | 0.095x |
| 28 | CES 210094 | 9532051 | 2.88 | 0.0894x |
| 29 | CES 210095 | 9370896 | 2.83 | 0.098x |
| 30 | CES 210096 | 9459180 | 2.86 | 0.0938x |
| 31 | CES 210097 | 8085352 | 2.44 | 0.0792x |

**Table S2: Final filtered distribution data for *T. truculentus* used for niche modelling**

| species | longitude | latitude |
| --- | --- | --- |
| *T_truculentus* | 76.01804 | 11.59391 |
| *T_truculentus* | 75.9436 | 11.86135 |
| *T_truculentus* | 75.57553 | 12.21464 |
| *T_truculentus* | 75.75021 | 12.41093 |
| *T_truculentus* | 75.57756 | 12.41253 |
| *T_truculentus* | 75.68222 | 12.84444 |
| *T_truculentus* | 75.42509 | 13.15268 |
| *T_truculentus* | 75.13171 | 13.27544 |
| *T_truculentus* | 75.08872 | 13.5155 |
| *T_truculentus* | 75.74972 | 13.53361 |
| *T_truculentus* | 75.18406 | 13.59146 |
| *T_truculentus* | 74.66004 | 14.25918 |
| *T_truculentus* | 74.64654 | 14.6426 |
| *T_truculentus* | 74.9405 | 14.68935 |
| *T_truculentus* | 74.47258 | 14.73427 |
| *T_truculentus* | 74.70738 | 14.75187 |
| *T_truculentus* | 74.84989 | 14.77363 |
| *T_truculentus* | 74.13232 | 14.82035 |
| *T_truculentus* | 74.59834 | 15.13944 |
| *T_truculentus* | 74.6003 | 15.24557 |
| *T_truculentus* | 74.05067 | 15.33291 |
| *T_truculentus* | 74.72932 | 15.35601 |
| *T_truculentus* | 75.7442 | 15.37231 |
| *T_truculentus* | 75.14425 | 15.3755 |
| *T_truculentus* | 74.14497 | 15.39267 |
| *T_truculentus* | 74.32639 | 15.41694 |
| *T_truculentus* | 74.37833 | 15.51556 |
| *T_truculentus* | 74.01608 | 15.72993 |
| *T_truculentus* | 74.17427 | 15.80556 |
| *T_truculentus* | 74.06274 | 15.96122 |
| *T_truculentus* | 74.20889 | 16.04861 |
